# Neuronal wiring diagram of an adult brain

**DOI:** 10.1101/2023.06.27.546656

**Authors:** Sven Dorkenwald, Arie Matsliah, Amy R Sterling, Philipp Schlegel, Szi-chieh Yu, Claire E. McKellar, Albert Lin, Marta Costa, Katharina Eichler, Yijie Yin, Will Silversmith, Casey Schneider-Mizell, Chris S. Jordan, Derrick Brittain, Akhilesh Halageri, Kai Kuehner, Oluwaseun Ogedengbe, Ryan Morey, Jay Gager, Krzysztof Kruk, Eric Perlman, Runzhe Yang, David Deutsch, Doug Bland, Marissa Sorek, Ran Lu, Thomas Macrina, Kisuk Lee, J. Alexander Bae, Shang Mu, Barak Nehoran, Eric Mitchell, Sergiy Popovych, Jingpeng Wu, Zhen Jia, Manuel Castro, Nico Kemnitz, Dodam Ih, Alexander Shakeel Bates, Nils Eckstein, Jan Funke, Forrest Collman, Davi D. Bock, Gregory S.X.E. Jefferis, H. Sebastian Seung, Mala Murthy, the FlyWire Consortium

## Abstract

Connections between neurons can be mapped by acquiring and analyzing electron microscopic (EM) brain images. In recent years, this approach has been applied to chunks of brains to reconstruct local connectivity maps that are highly informative, yet inadequate for understanding brain function more globally. Here, we present the first neuronal wiring diagram of a whole adult brain, containing 5×10^7^ chemical synapses between ∼130,000 neurons reconstructed from a female *Drosophila melanogaster*. The resource also incorporates annotations of cell classes and types, nerves, hemilineages, and predictions of neurotransmitter identities. Data products are available by download, programmatic access, and interactive browsing and made interoperable with other fly data resources. We show how to derive a projectome, a map of projections between regions, from the connectome. We demonstrate the tracing of synaptic pathways and the analysis of information flow from inputs (sensory and ascending neurons) to outputs (motor, endocrine, and descending neurons), across both hemispheres, and between the central brain and the optic lobes. Tracing from a subset of photoreceptors all the way to descending motor pathways illustrates how structure can uncover putative circuit mechanisms underlying sensorimotor behaviors. The technologies and open ecosystem of the FlyWire Consortium set the stage for future large-scale connectome projects in other species.

## Nomenclature

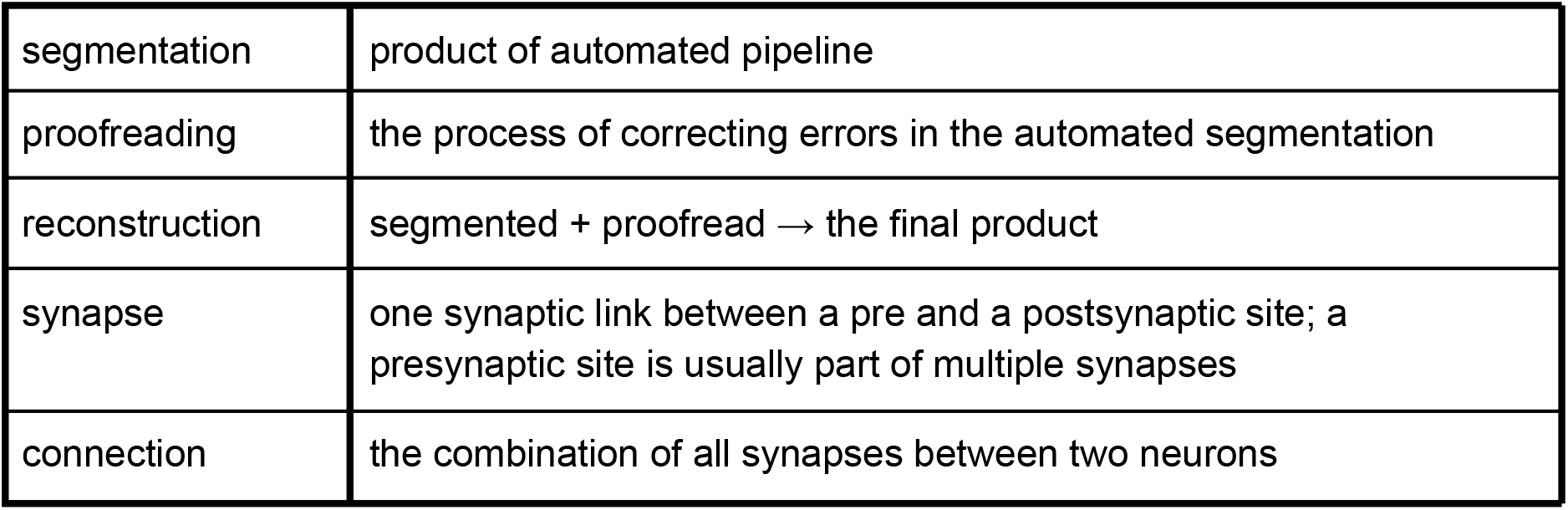

|  |  |
| --- | --- |
| segmentation | product of automated pipeline |
| proofreading | the process of correcting errors in the automated segmentation |
| reconstruction | segmented + proofread → the final product |
| synapse | one synaptic link between a pre and a postsynaptic site; a presynaptic site is usually part of multiple synapses |
| connection | the combination of all synapses between two neurons |

## Introduction

While rudimentary nervous systems existed in more ancient animals^1^, brains evolved perhaps half a billion years ago^2^, and are essential for the generation of sophisticated behaviors. It is widely accepted that dividing a brain into regions is helpful for understanding brain function^3^. Wiring diagrams at the level of neurons and synapses have been controversial^4–6^. Skepticism flourished largely due to a lack of technologies that could reconstruct such wiring diagrams^7,8^. The situation began to change in the 2000s, due to the efforts of a small community of researchers. Here we report a significant milestone attained by these efforts, the first neuronal wiring diagram of a whole adult brain.

The brain of *Drosophila melanogaster* may seem tiny, but its 10^5^ neurons and 10^8^ synapses enable a fly to see, smell, hear, walk, and, of course, fly. Flies engage in dynamic social interactions^9^, navigate over distances^10^, and form long-term memories^11^. Portions of fly brains have been reconstructed from electron microscopic (EM) images, which have sufficient resolution to reveal the fine branches of neurons and the synapses that connect them. The resulting wiring diagrams of neural circuits have provided crucial insights into how the brain generates social^12,13^, memory-related^14^ or navigation^15^ behaviors. Wiring diagrams of other fly brain regions have been mapped and related to visual^16,17^, auditory^18^, and olfactory^14,19,20^ functions. Similarities with mammalian wiring diagrams ^21–23^ are striking.

The above wiring diagrams and many others from mammals^24–28^ have come from pieces of brain. But recordings of *Drosophila* neural activity have revealed nearly brain-wide encoding of sensory^29^ and motor^30–32^ variables. These studies and others in vertebrates highlight that understanding how the brain processes sensory information or drives behavior will require understanding global information flow at the scale of the entire brain.

The closest antecedent to our whole brain is the reconstruction of a fly “hemibrain”^33^, a pioneering resource that has already become indispensable to *Drosophila* researchers^14,20,34,35^. It is estimated to contain about 20,000 neurons that are “uncropped,” i.e., minimally truncated by the borders of the imaged volume, and 14 million synapses between them. Our reconstruction of an entire adult brain contains 127,978 neurons (Fig. 1a), and 53 million synapses between them. These and many other data products (Fig. 1b) are available for download, programmatic access, and interactive browsing and made interoperable with other fly data resources through a growing ecosystem of software tools (Fig. 1c). The primary portal to the data is FlyWire Codex (codex.flywire.ai, manuscript *in prep*), which makes the information visualizable and queryable.

**Figure 1.**
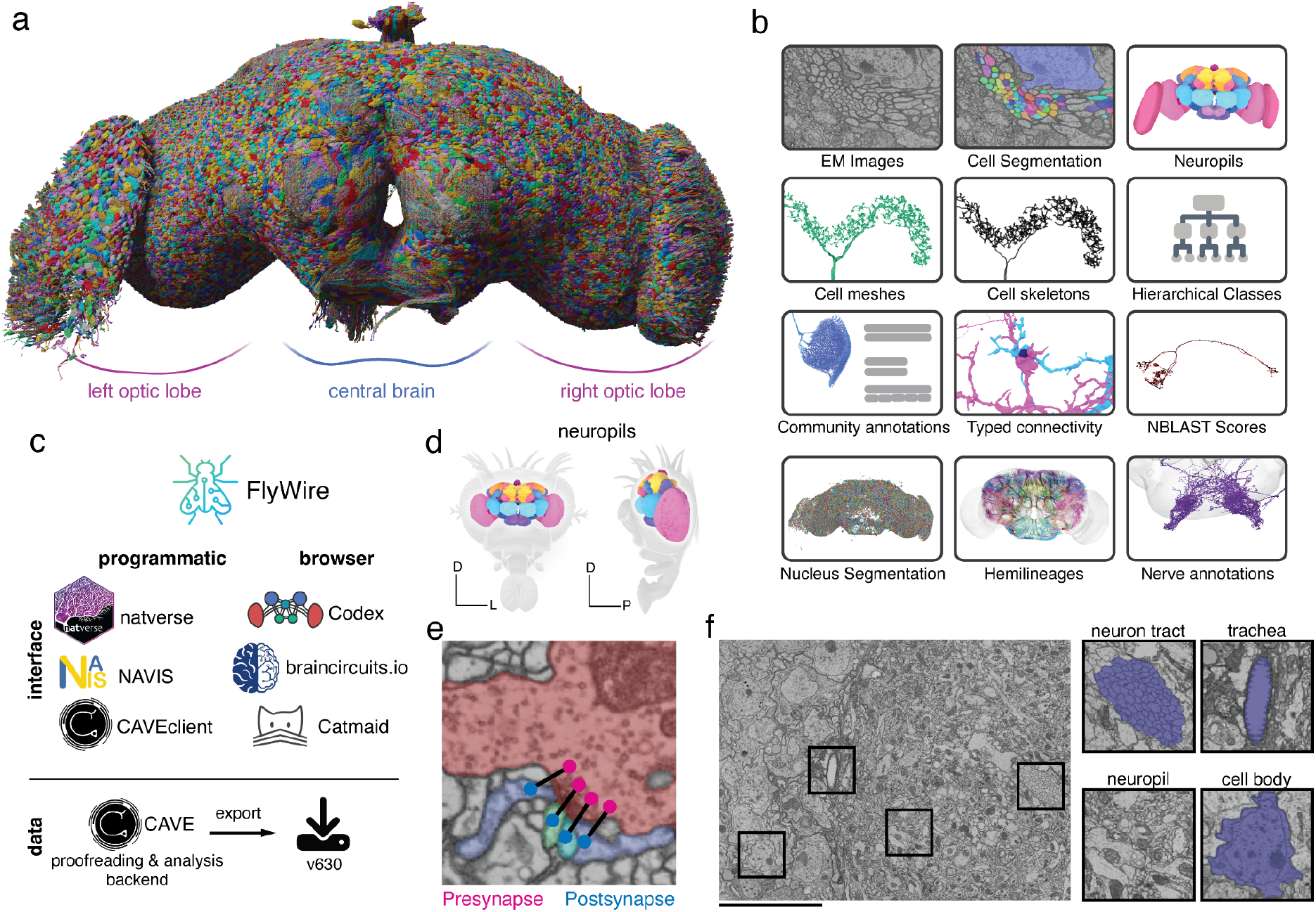
A connectomic reconstruction of a whole fly brain. (a) All neuron morphologies reconstructed with FlyWire. All neurons in the central brain and both optic lobes were segmented and proofread. Note: image and dataset are mirror inverted relative to the native fly brain. (b) An overview of many of the FlyWire resources which are being made available. FlyWire leverages existing resources for EM imagery by Zheng et al.^52^, synapse predictions by Buhmann et al.^45,46^ and neurotransmitter predictions by Eckstein et al.^47^. Annotations of the FlyWire dataset such as hemilineages, nerves, and hierarchical classes are established in our companion paper by Schlegel et al. (c) FlyWire uses CAVE (*in prep*) for proofreading, data management, and analysis backend. The data can be accessed programmatically through the CAVEclient, navis and natverse^168^, and through the browser in Codex, Catmaid Spaces and braincircuits.io. Static exports of the data are also available. (d) The *Drosophila* brain can be divided into spatially defined regions based on neuropils^110^ (Ext. Data Fig. 1-1). Neuropils for the lamina are not shown. (e) Synaptic boutons in the fly brain are often polyadic such that there are multiple postsynaptic partners per presynaptic bouton. Each link between a pre- and a postsynaptic location is a synapse. (f) Neuron tracts, trachea, neuropil, cell bodies can be readily identified from the EM data which was acquired by Zheng et al.^52^. Scale bar: 10 μm

The wiring diagram from our whole brain reconstruction is complete enough to deserve the name “connectome.” It is a clear leap beyond *C. elegans* (300 neurons, <10^4^ synapses)^36–38^ and the 1st instar larva of *Drosophila* (3,000 neurons, 5×10^5^ synapses)^39^. Our connectome advances beyond the hemibrain in ways that are not simply numerical. It encompasses the subesophageal zone (SEZ) of the central brain, important for diverse functions such as gustation and mechanosensation (see companion paper Shiu et al.^40^ as well as Eichler et al.^41^), and containing many of the processes of neurons that descend from the brain to the ventral nerve cord to drive motor behaviors. It includes annotations for nearly all sexually-dimorphic neurons, analyzed in a companion paper (Deutsch et al., *in prep*). Our reconstruction of both optic lobes goes far beyond existing maps of columnar visual circuitry^17,42,43^. Connections between the optic lobes and central brain are included, as explored by a companion paper (Kind, Garner et al., *in prep*). Also included are neurons that extend into the brain through the nerves and neck connective, which are essential for tracing sensorimotor pathways, as illustrated by the present paper and companion papers.

Our reconstruction utilized image acquisition and analysis techniques that are distinct from those used for the hemibrain (Methods and Discussion). However, we have built directly on the hemibrain in an important way. The companion paper by Schlegel et al. annotated cell types of central brain neurons, principally by matching them with hemibrain neurons. This approach was enabled by a growing ecosystem of software tools serving interoperability between different fly data sources (Fig. 1c). Because annotations of cell types are essential for scientific discovery, Schlegel et al.^44^ should be cited along with the present manuscript by those who use the FlyWire resource. Annotations in the SEZ and optic lobes, largely absent from the hemibrain, were contributed by *Drosophila* labs in the FlyWire Consortium, private corporations, and citizen scientists. Synapse predictions^45,46^ and estimates of neurotransmitter identities^47^ were also contributed by the community.

After matching, Schlegel et al.^44^ have also compared our wiring diagram with the hemibrain where they overlap and showed that cell type counts and large strong connections were largely in agreement. This means that the combined effects of natural variability across individuals and “noise” due to imperfect reconstruction tend to be modest, so our wiring diagram of a single brain should be useful for studying any normal *Drosophila* individual. That being said, there are known male-female differences^48^. In addition, our companion paper reports high variability for principal neurons of the mushroom body, a brain structure required for olfactory learning and memory^44^. Some mushroom body connectivity patterns have even been found to be near random^49,50^, though deviations from randomness have since been identified^51^. In short, *Drosophila* wiring diagrams are useful because of their stereotypy, yet also open the door to studies of connectome variation.

In addition to describing the FlyWire resource, this manuscript also presents analyses that illustrate how the data products can be used. Additional whole-brain network analyses are provided in a companion paper (Lin et al., *in prep*). From the connectome with its huge numbers of neurons and synapses, we derive a projectome, a reduced map of projections between 78 fly brain regions known as neuropils (Fig. 1d, Ext. Data Fig. 1-1). We trace synaptic pathways and analyze information flow from the inputs to the outputs of the brain, across both hemispheres, and between the central brain and the optic lobes. In particular, the organization of excitation and inhibition in pathways from photoreceptors in the ocelli to descending motor neurons immediately suggests hypotheses about circuit mechanisms of behavior.

## Results

### Reconstruction of a whole fly brain at electron microscopic resolution

Images of an entire adult female fly brain (Fig. 1e, f) were previously acquired by serial section transmission EM, and released into the public domain by Zheng et al.^52^. We previously realigned the EM images^53^, automatically segmented all neurons in the images^54^, created a computational system that allows interactive proofreading of the segmentation, and assembled an online community known as FlyWire^55^. During the initial phase, much proofreading was done by a distributed community of *Drosophila* labs in the FlyWire Consortium, and focused on neurons of interest to these labs. During the later phase, the remaining neurons were mainly proofread by two centralized teams at Princeton and Cambridge, with significant contributions from citizen scientists worldwide. The recruitment and training of proofreaders and their workflows are described in the Methods.

Chemical synapses were automatically detected in the images as pairs of presynapse-postsynapse locations^45,46^. The whole brain contains 0.0188 mm^3^ of neuropil volume and ∼130 million synapses. This works out to 6.9 synapses/µm^3^, much denser than the <1 synapse/µm^3^ reported for mammalian cortex^56,57^. The central brain and left and right optic lobes contain 0.0103, 0.0042, and 0.0043 mm^3^ of neuropil volume, respectively, with synapse counts in approximately the same proportion. Synapses were combined with proofread neurons to yield the connectome, using the Connectome Annotation Versioning Engine (CAVE*, in prep*).

We already showed that FlyWire proofreading can yield accurate results^55^ through comparison with light microscopic reconstructions of neurons that are known to be highly stereotyped across individual flies. A second method is to subject neurons to an additional round of proofreading^33,58^, which was previously shown to yield few changes^55^. Because proofreading workflows and personnel have changed over time, and accuracy can vary across brain regions, we repeated this evaluation by subjecting 826 neurons from the central brain to a second round of proofreading. Relative to the second round, our first round of proofreading achieved an average F1-Score of 99.2% by volume (Ext. Data Fig. 1-2 a,b).

A third validation method is to quantify how many of the automatically detected synapses are attached to proofread segments, as opposed to being isolated in tiny “orphan” segments^45,46^. We found high attachment rates of presynapses (92.3% or ∼120,100,000 presynapses attached) while attachment rates of postsynapses were lower (43.9% or ∼57,200,000 postsynapses attached) due to less proofreading and reattachment of twigs which contain most of the postsynapses^55^ (Ext. Data Fig. 1-2 c,d). Attachment rates were generally in agreement between the two hemispheres of FlyWire and with the hemibrain (Ext. Data Fig. 1-2 e,f,g) and varied by neuropil (Ext. Data Fig. 1-3). The bottom line is that accuracy of our connectome is state-of-the-art. As with the hemibrain^33^, false negative synapses are the dominant kind of error but false positives exist as well. For this reason all analyses we present below (and connections indicated in Codex) use a threshold of 5 synapses to determine a connection between two neurons. Assuming that such errors are statistically independent, accuracy is expected to be high for detection of connections involving multiple synapses^33,44,59,60^.

FlyWire’s reconstruction remains open for proofreading and annotations and new versions of the resource will be released in future. This allows for the correction of remaining errors as they are discovered and further rounds of validation to be performed. Additionally, as explained below, proofreading of photoreceptor axons in the compound eyes is still ongoing. The first public release (called version 630) has been extensively validated for neurons in the central brain. All neurons in the optic lobe were proofread but additional validation will likely identify and correct minor reconstruction errors.

### Intrinsic neurons of the brain

Of the 127,978 proofread neurons in FlyWire, 114,423 are fully contained within the brain (including both central brain and optic lobes, but excluding afferent and efferent neurons, with projections into and out of the brain, respectively; Fig. 2a,b). These intrinsic neurons (Fig. 2c left) belong to the brain only, in contrast to other neurons that are shared by the brain with other structures. Intrinsic neurons of the brain make up three quarters of the adult fly nervous system (Methods), indicating a high degree of centralization in the brain. The large fraction is related to the fact that in the adult the brain is substantially larger than the ventral nerve cord (VNC)^61–63^. Intrinsic neurons amount to 84% of brain neurons. Their predominance means that the brain primarily communicates with itself, and only secondarily with the outside world.

**Figure 2.**
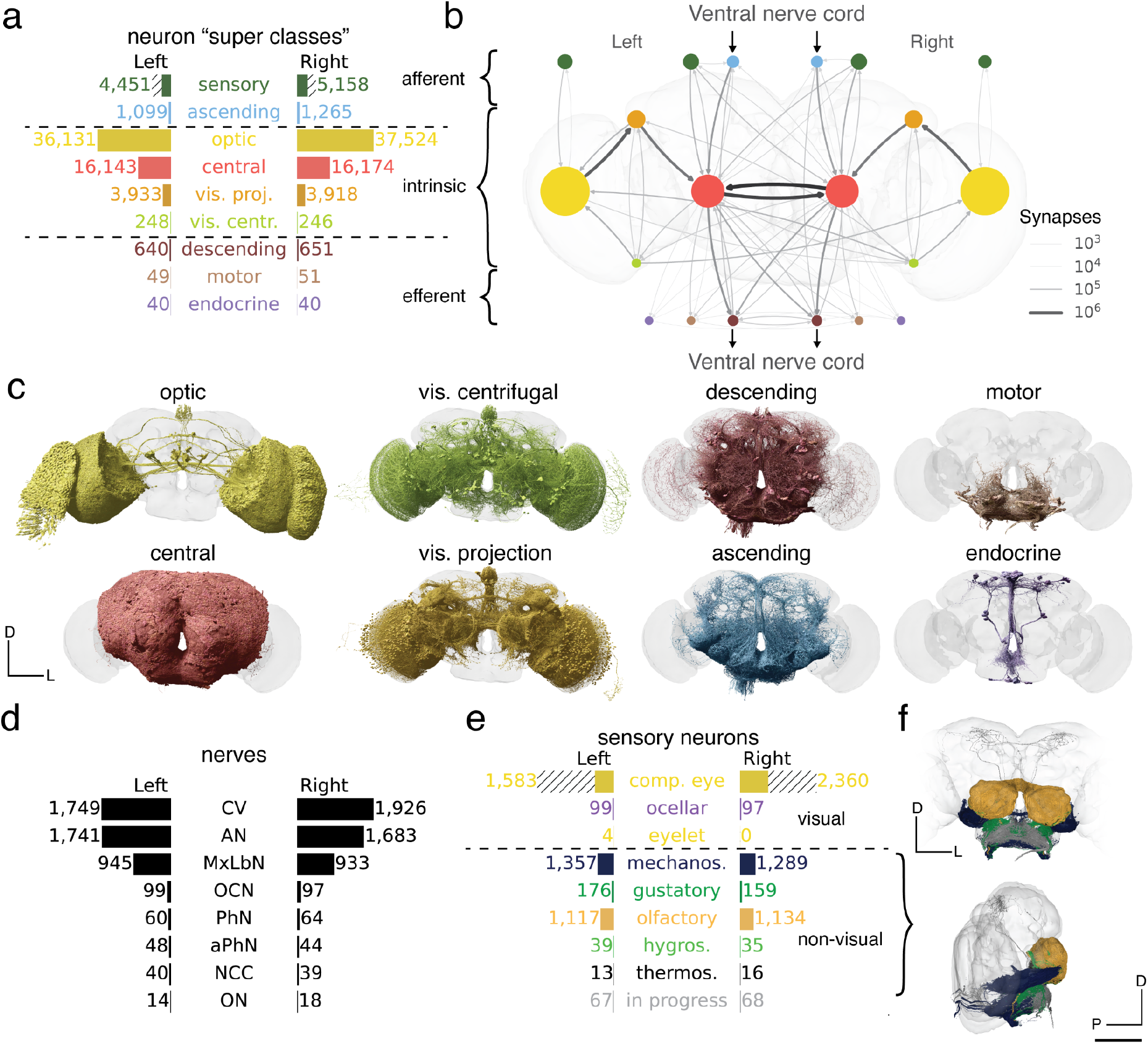
Neuron categories. (a) We grouped neurons in the fly brain by “flow”: intrinsic, afferent, efferent. Each flow class is further divided into “super-classes’’ based on location and function. Neuron annotations are described in more detail in our companion paper by ^44^. The first public release is missing ∼8,000 retinula cells in the compound eyes and four eyelets in one hemisphere which are indicated by hatched bars. (b) Using these neuron annotations, we created an aggregated synapse graph between the super-classes in the fly brain. (c) Renderings of all neurons in each super-class. (d) There are eight nerves into each hemisphere in addition to the ocellar nerve and the cervical connective (CV). All neurons traversing the nerves have been reconstructed and accounted for. (e) Sensory neurons can be subdivided by the sensory modality they respond to. In FlyWire, almost all sensory neurons have been typed by modality. The counts for the medial ocelli were omitted and are shown in Fig. 7b. (f) Renderings of all non-visual sensory neurons. Scale bar: 100 µm

The nervous system of the larval fly is less centralized; intrinsic neurons of the brain make up one quarter to one third of its nervous system^39^. The closest structure to a brain in *C. elegans* is the nerve ring^64^, which is co-located with multiple sensory organs in the worm’s head. The nerve ring contains no intrinsic neurons, as all neurons in the nerve ring also extend neurites into the rest of the nervous system. The absence of intrinsic neurons is consistent with the convention that the nerve ring is not commonly called a brain.

While the above statistics are based on neuron numbers, they are conceptually related to volume-based measures of encephalization used in studies of brain evolution^65^. For comparison, the rat brain occupies 65% of its central nervous system by volume^66^. Our neuron-based measure of encephalization cannot yet be computed for rodents, but this will become possible as connectomics continues to scale^67^.

### Afferent and efferent neurons

Brain neurons that are not intrinsic can be divided into two categories, depending on the locations of their cell bodies. For afferent (sensory, ascending) neurons, the cell body is outside the brain, while for efferent (descending, motor, endocrine) neurons, the cell body is contained in the brain. It is generally accurate to think of an afferent neuron as a brain input, and an efferent neuron as a brain output. The relation to information flow is actually more subtle, however, as most fly neurites carry some mixture of presynapses and postsynapses on both dendrites and axons^39,47,59,60^.

Our companion paper exhaustively identified all afferent and efferent neurons contained in cross sections of nerves and the neck connective running between the brain and VNC (Fig. 2d)^44^. Almost 95% of these neurons were in the neck connective, antennal nerve, and maxillary-labial nerve. Although afferents are truncated in our reconstruction, Schlegel et al.^44^ along with other community members^41,68^ were able to determine the sensory organs corresponding to 5,362 of the 5,495 non-visual sensory neurons (Fig. 2e,f). Non-visual sensory neurons enter the brain through nerves (Fig. 2d) that mostly terminate in the antennal lobe or the SEZ (we define the SEZ as containing the following neuropils: SAD, GNG, AMMC, and PRW^69^; see Ext. Data Fig. 1-1 for neuropil definitions). The antennal lobe (AL) is the first relay center for processing of olfactory information, and many of the olfactory receptor neuron (ORN) inputs to the AL were reconstructed in the hemibrain as well. The SEZ receives more diverse inputs, including the projections of both mechanoreceptor and gustatory receptor neurons - these projections were not contained in the hemibrain. The nerves contained few efferent neurons, among which were head motor neurons (N=100) or endocrine neurons (N=80) (Fig. 2a,b,c). A large fraction of efferent neurons have branches in the SEZ, including most of the 100 motor neurons.

Visual afferents are by far the most numerous kind of sensory input, and enter the brain directly rather than through nerves. This is the last class of neuron that remains to be fully proofread. There are photoreceptor axons coming from the compound eyes (∼12,800, of which 3,943 have already been proofread in both eyes), ocelli (270 of which all have been proofread), and eyelets (8 of which 4 have been proofread).

The neurons traversing the neck connective were grouped into 1,303 efferent (descending) and 2,364 afferent (ascending) neurons (Fig. 2a,b,c). In a companion paper, Eichler et al. (*in prep*) typed these neurons and matched them to reconstructions from two separate EM datasets of a VNC^61,70,71^, allowing circuits spanning the whole CNS (brain and VNC) to be at least schematically mapped.

### Optic lobes and central brain

Of the 114,423 intrinsic neurons, 32,422 are fully contained in the central brain, and 73,655 are fully contained in the optic lobes and ocellar ganglia (this number excludes the photoreceptors, which are sensory afferent neurons, see above). Given that the visual areas dominate the count, it seems safe to say that *Drosophila* is a highly visual animal. The optic lobes, which are largely absent from the 1st instar larval, are a major reason that the adult fly brain so dominates its nervous system.

The optic lobes and ocellar ganglia also contain 7,851 neurons that project into the central brain, so called visual projection neurons (VPNs)^44^. We provide a more detailed analysis of connections in the ocellar ganglion in Fig. 7. Many VPNs are columnar types that tile the visual field. VPNs target specific neuropils (e.g., AOTU, PLP, and PVLP) or optic glomeruli^72,73^ in the central brain. The influence of VPNs can be very strong; 879 central neurons receive more than half their synapses from VPNs.

The hemibrain already characterized several VPN types along with their outputs in the central brain^33^. Our whole brain reconstruction reveals many other aspects of VPN connectivity, such as their inputs in the medulla, lobula, and lobula plate. In addition to feedforward targeting of central neurons, VPNs make 20% of their synapses onto other VPNs, and 21% onto optic lobe neurons. Companion papers investigate the visual projections to the central complex and the mushroom body (Heckman and Clowney, *in prep*).

There are 494 neurons that project from the central brain to the optic lobes^44^. We call these visual centrifugal neurons (VCNs)^73^. They are distinct from previously defined types of visual centrifugal neurons that are fully contained in the optic lobe, and their functions are mostly unknown. VCNs are 15× less numerous than VPNs. Nevertheless, half of all optic lobe neurons receive 5 or more synapses from VCNs, showing that much early visual processing incorporates feedback from the central brain. Centrifugal inputs to the retina are found in many vertebrate species, including humans^74^.

Many VCNs arborize broadly in the optic lobe, appearing to cover the entire visual field. Some VCNs, however, cover only a subset of columns within a portion of the visual field. A few optic lobe neurons receive as many as 50% of their synapses from VCNs. These belong to the class of peptidergic neurons involved in circadian rhythmicity, which are detailed in a companion paper (Reinhard and Fukuda et al. et al., *in prep*). Tm5c is a columnar type (necessary for *Drosophila*’s preference for UV over visible light^75^) with more than 10% of its input from VCNs.

A lamina wide-field neuron (Lawf2) can receive more than 10% of its input from VCNs, and a major input source is octopaminergic (OA-AL2b2). It was previously shown that gain modulation of Lawf2 neurons increases during flight^76^, and this effect is mimicked by bath application of octopamine. Transcriptomic studies showed that Lawf2 neurons express octopamine receptors at high levels^77^.

### Neuron super-classes

The neuron classes introduced above are organized into a hierarchy, as explained in our companion paper^44^. The three “flow” classes (afferent, intrinsic, efferent) are divided into the nine “super-classes” mentioned above (Fig. 2a). A simplified representation of the connectome as a graph in which nodes are super-classes is shown in Fig. 2b. Node sizes reflect neuron number, and link widths indicate connection number. This is the first of several simplified representations that we will introduce to tame the complexity of the connectome.

### Neurons and glia

A basic property of the fly brain is that cell bodies are spatially segregated from neurites. Cell bodies reside near the surface (“rind”) of the brain (Fig. 3a), surrounding a synapse-rich interior that mainly consists of entangled neurons and glia, fiber bundles or tracts, as well as tubules of the tracheal system (Fig. 1f, Ext. Data Fig. 1-4a, Colodner et al*, in prep*).

**Figure 3.**
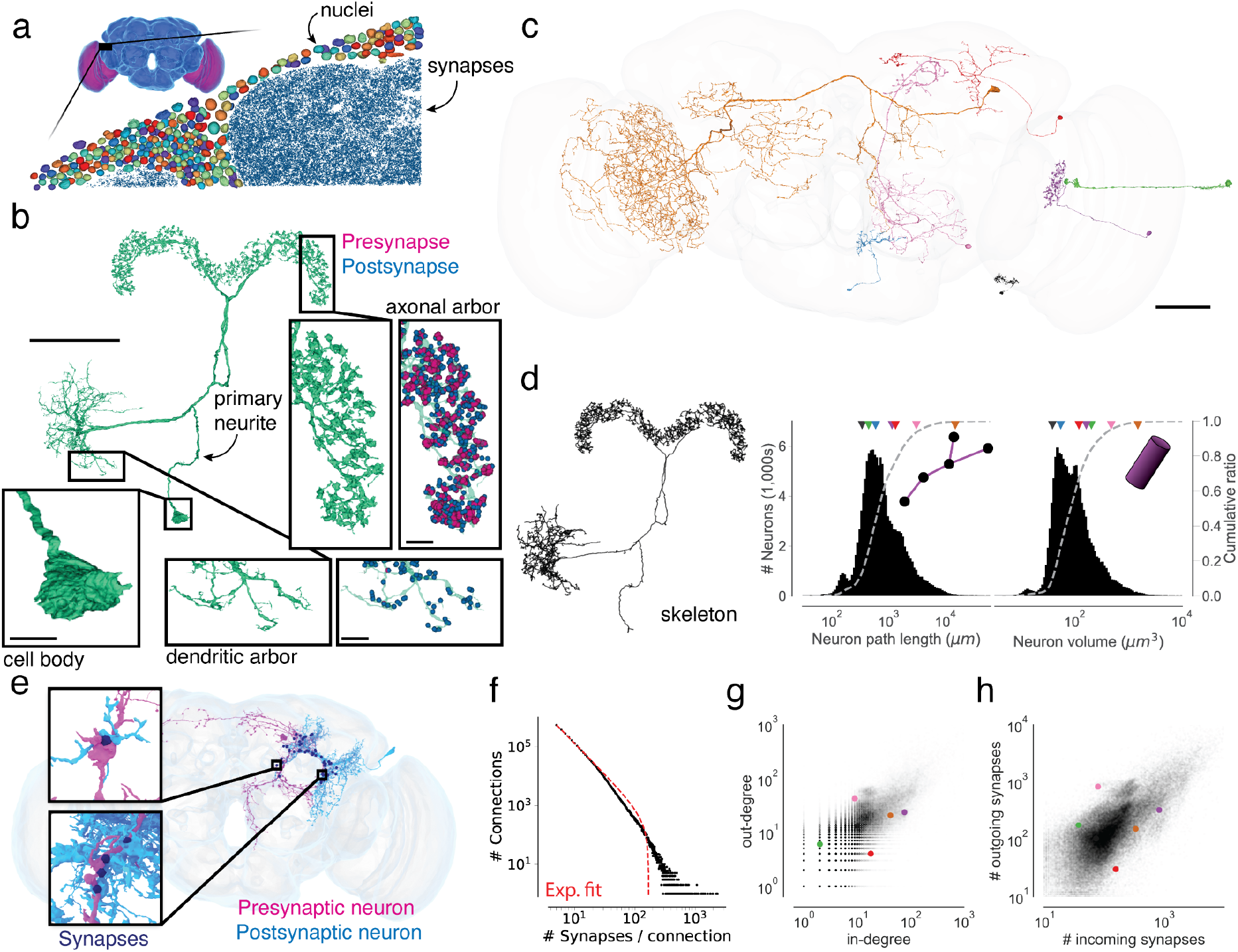
Neuron and connection sizes. (a) The synapse-rich (synapses in blue) neuropil is surrounded by a layer of nuclei (random colors) located at the outside of the brain as well as between the optic lobes (purple) and the central brain (blue). (b) An LPsP neuron can be divided into morphologically distinct regions. Synapses (purple and blue) are found on the neuronal twigs and only rarely on the backbone. (c) We selected seven diverse neurons as a reference for the following panels. (d) The morphology of a neuron can be reduced to a skeleton from which the path length can be measured. The histograms show the distribution of path length and volume (the sum of all internal voxels) for all neurons. The triangles on top of the distributions indicate the measurements of the neurons in (b). (e) Connections in the fly brain are usually multisynaptic as in this example of neurons connecting with 71 synapses. (f) The number of connections with a given number of synapses and a fitted truncated power law distribution. (g) In degree and out degree of intrinsic neurons in the fly brain are linearly correlated (R=0.76). (h) The number of synapses per neuron varies between neurons by over a magnitude and the number of incoming and outgoing synapses is linearly correlated (R=0.80). Only intrinsic neurons were included in this plot. Scale bars: 50 μm (b, c), 10 µm (b-insets)

A typical non-sensory *Drosophila* neuron is unipolar and consists of a primary neurite (also known as cell body fiber) that leaves the cell body (soma), enters the neuropil, and branches into secondary and higher-order neurites (Fig. 3b). Secondary neurites can sometimes be classified as axons if presynapses clearly dominate, or dendrites if postsynapses clearly dominate^39,47,59^. Such an axon-dendrite distinction was made, for example, when defining visual projection and centrifugal neurons above.

But some mixture of presynapses and postsynapses is generally found on all non-primary neurites^39,47,59,60^ (Fig. 3b). In addition, the soma of insect neurons is separated from the main processes (Fig. 3b). Given this structure, the concept that signals pass from dendrites to soma to axon, which is often a good approximation for mammalian neurons, may not apply for non-sensory neurons in the fly.

Neurons vary greatly in size and shape (Fig. 3c). We computed skeletons for all reconstructed neurons (Fig. 3d) to measure neuronal path lengths. The median path length of a neuronal arbor was 656 µm (Fig. 3d). It has been argued that branched arbors are optimal for achieving a high degree of connectivity with other neurons^78^. Neurons with short path lengths are interesting exceptions, and can be found in both the optic lobes and central brain. Path length and volume both varied over two orders of magnitude (Fig. 3d, path length percentiles: 0.1%: 0.059 mm, 99.9%: 19.211 mm, volume percentiles: 0.1%: 80 µm^3^, 99.9%: 459 µm^3^). In total, the brain contains ∼146 m of neuronal path length.

Sizes vary significantly between different cell super-classes (Ext. Data Fig. 3-1a-f). Optic lobe neurons are on average much shorter than central brain neurons (0.70 mm vs 2.15 mm on average) and take up a smaller volume (0.0066 mm^3^ vs 0.0086 mm^3^ total neuronal volume), which is why the optic lobes dominate the brain by neuron number but not by volume or synapse count. Visual centrifugal neurons are among the largest in the brain, and larger on average than visual projection neurons (5.05 mm vs 1.56 mm on average). While we measured much shorter path lengths and volumes for afferent neurons because only part of their axonal arbors is contained within the brain (Ext. Data Fig. 3-1b,e), arbors of efferents, motor and descending neurons which also have some of their arbor outside the brain, were among the largest we measured (Ext. Data Fig. 3-1c,f).

A small fraction of brain volume is glial cells, which are categorized in six types^79,80^. We estimated that 13% of the cell bodies in the EM dataset are non-neuronal or glial^81^. Only a few astrocyte-like glia have been proofread (Ext. Data Fig. 1-4b). Sheet-like fragments of ensheathing glia are readily found near fiber bundles in the automated reconstruction. Further proofreading of glia could be prioritized in the future if there is community demand.

### Synapses and connections

Our connectome includes only chemical synapses; the identification of electrical synapses awaits a future EM dataset with higher resolution (see Discussion). We use the term “synapse” to mean chemical synapse. A *Drosophila* synapse is generally polyadic, meaning that a single presynapse communicates with multiple target postsynapses (Fig. 1e). FlyWire represents a polyadic synapse as multiple synapses, each of which is a pair of presynaptic and postsynaptic locations^45^. Polyadic synapses are common in other invertebrate species, such as *C. elegans*, and exist in some mammalian brain structures (e.g. retina).

We define a connection from neuron A to neuron B as the set of synapses from A to B. A connection typically contains multiple synapses, and the number can be large (Fig. 3 e,f). Connections with less than 10 synapses are typical, but a single connection can comprise >100 synapses (N=14,969) or even >1,000 synapses (N=27). The strongest connection was from a visual centrifugal neuron (LT39) onto a wide field lobula neuron (mALC2), and contained over 2300 synapses.

These numbers are much larger than the report of a maximum of 41 synapses connecting a pair of *C. elegans* neurons^38^. To model such a distribution with a long tail, we used a power law with exponential cutoff^33^ (Fig. 3g). Our fit found comparable parameters, but the fit to our whole-brain distribution of connection strengths was not as good as their fit to the hemibrain distribution. A similar power law is also a reasonable fit to the distribution of connection strengths in *C. elegans*.

Setting a threshold of ≥5 synapses for determining a (strong) connection is likely to be adequate for avoiding false positives in the dataset, but not missing connections (see Methods). There are 2,613,129 such connections between the 124,891 identified neurons. There are several reasons to focus on strong connections. First, a connection with many synapses is expected to be strong in a physiological sense, other things being equal^82–84^. Second, strong connections are likely to be more reproducible across individuals^44,85,86^. Third, higher accuracy (both precision and recall) of automatic detection is expected for strong connections, assuming that errors are statistically independent^33,59^.

One of the most basic properties of a node in any network is its degree, the number of nodes to which it is linked. To characterize the degree distribution in the *Drosophila* connectome, we focused on intrinsic neurons (N=114,423) because, unlike afferent and efferent neurons, they do not suffer from undercounting of connections due to truncation.

For any neuron, in-degree is defined as its number of presynaptic partners (input neurons), and out-degree is defined as its number of postsynaptic partners (output neurons). The median in-degree and out-degree of intrinsic neurons are 11 and 13 (Fig. 3g), respectively, with the restriction mentioned above to connections involving five or more synapses. These median values do not seem dramatically different from the median in-degree and out-degree of 10 and 19 for neurons in the *C. elegans* hermaphrodite, considering that the latter contains several hundred times fewer neurons than *Drosophila*.

The neuron in the *Drosophila* brain with maximum degree is a visual GABAergic interneuron (CT1), with 6329 postsynaptic partners and 4999 presynaptic partners. CT1 arborizes exclusively in the medulla neuropil of the optic lobe - indeed, most neuropils of the *Drosophila* brain contain one or a few large GABAergic neurons private to that neuropil, with high in-degree and out-degree (see Lin et al., *in prep*, for more analysis on connectivity motifs in FlyWire); these neurons are considered to be important for local feedback gain control^87,88^. In a *C. elegans* hermaphrodite^38^, the neuron with maximum degree is a command interneuron for backward locomotion (AVAL), with 110 postsynaptic partners and 64 presynaptic partners. The existence of neurons with much higher degree is a marked way in which the *Drosophila* connectome differs from that of *C. elegans*. That being said, the degree of AVAL is large in a relative sense because it is a large fraction of the total *C. elegans* neuron number (302).

The number of synapses established by a neuron is correlated with its total neurite path length (R=0.80 (pre), R=0.89 (post), Ext. Data Fig. 3-1g). Presynapse and postsynapse counts are similarly correlated per neuron (R=0.80, Fig. 3h). We asked whether large neurons tend to use their many synapses to create stronger connections with individual neurons versus more connections with many different neurons. The total number of synapses established by a neuron was much better correlated with its in and out degrees (R=0.93, R=0.93 respectively) than its average connection strength (R=0.26, R=0.31 respectively, Ext. Data Fig. 3-1h,i). It remains to be tested whether the additional partners are from the same or different cell types.

Connections and neurons are not necessarily the functional units of neural computation. For certain large fly neurons, the arbors are composed of multiple compartments that function somewhat independently^89–91^. Perhaps these subcellular compartments, rather than whole cells, should be regarded as nodes of the connectome. Then CT1 would be replaced by many nodes with lower degrees. And the connection from LT39 to mALC2 would be replaced by many connections with fewer synapses between compartments of these neurons. A connectome of neuronal compartments can in principle be studied using our resource, which includes the location of every synapse.

### Neurotransmitter identity

A statistical prediction of the small molecule neurotransmitter (GABA, glutamate, acetylcholine, serotonin, dopamine, and octopamine) secreted by each neuron is available. A number of validations suggest that the predictions are highly accurate in aggregate^47^, though for any given synapse the prediction could be wrong. We assume that every neuron secretes a single small molecule neurotransmitter and combine the predictions for all outgoing synapses to an estimate which we assign to all outgoing synapses of a neuron, i.e. we provisionally assume neurons obey Dale’s law, although it is known that co-transmission does occur in the fly brain^92–95^.

GABAergic and glutamatergic neurons had much higher degrees than cholinergic neurons (Ext. Data Fig. 3-1j). Across all neuron categories, we found that GABAergic neurons were on average longer than glutamatergic and cholinergic neurons (Ext. Data Fig. 3-1k).

As a rule, we will assume that cholinergic neurons are excitatory and GABAergic and glutamatergic neurons are inhibitory^96–99^. A companion paper identifies all GABAergic and glutamatergic neurons that are bidirectionally coupled with large numbers of cholinergic neurons (Lin et al., *in prep*). This reciprocal inhibitory-excitatory motif is widespread throughout the fly brain^33,100^.

### From connectome to projectome

For mammals, tracer injection studies have mapped the axonal projections between brain regions of mouse^101–103^ and macaque^104,105^. In fly, large numbers of light microscopic reconstructions of single neurons have been aggregated to map projections between brain regions^106–108^. Such maps have been called projectomes^109^ or mesoscale connectomes^4^. In such techniques, the sampling of axons is difficult to control, which means that accurate quantification of projection strength is challenging.

Here we compute a projectome from a synapse-level connectome (Fig. 4a, Ext. Data Fig. 4-1). The interior of the fly brain has been subdivided into hierarchical neuropil regions^110^ (Ext. Fig. 1-1, Fig. 1d). Our fly projectome is defined as a map of projections between these neuropil regions. Because cell bodies are spatially separated from neuropils, a fly neuron cannot typically be assigned to a single brain region. This is unlike the situation for a mammalian neuron, which is conventionally assigned to the region containing its cell body. A typical fly neuron belongs to multiple neuropils.

**Figure 4.**
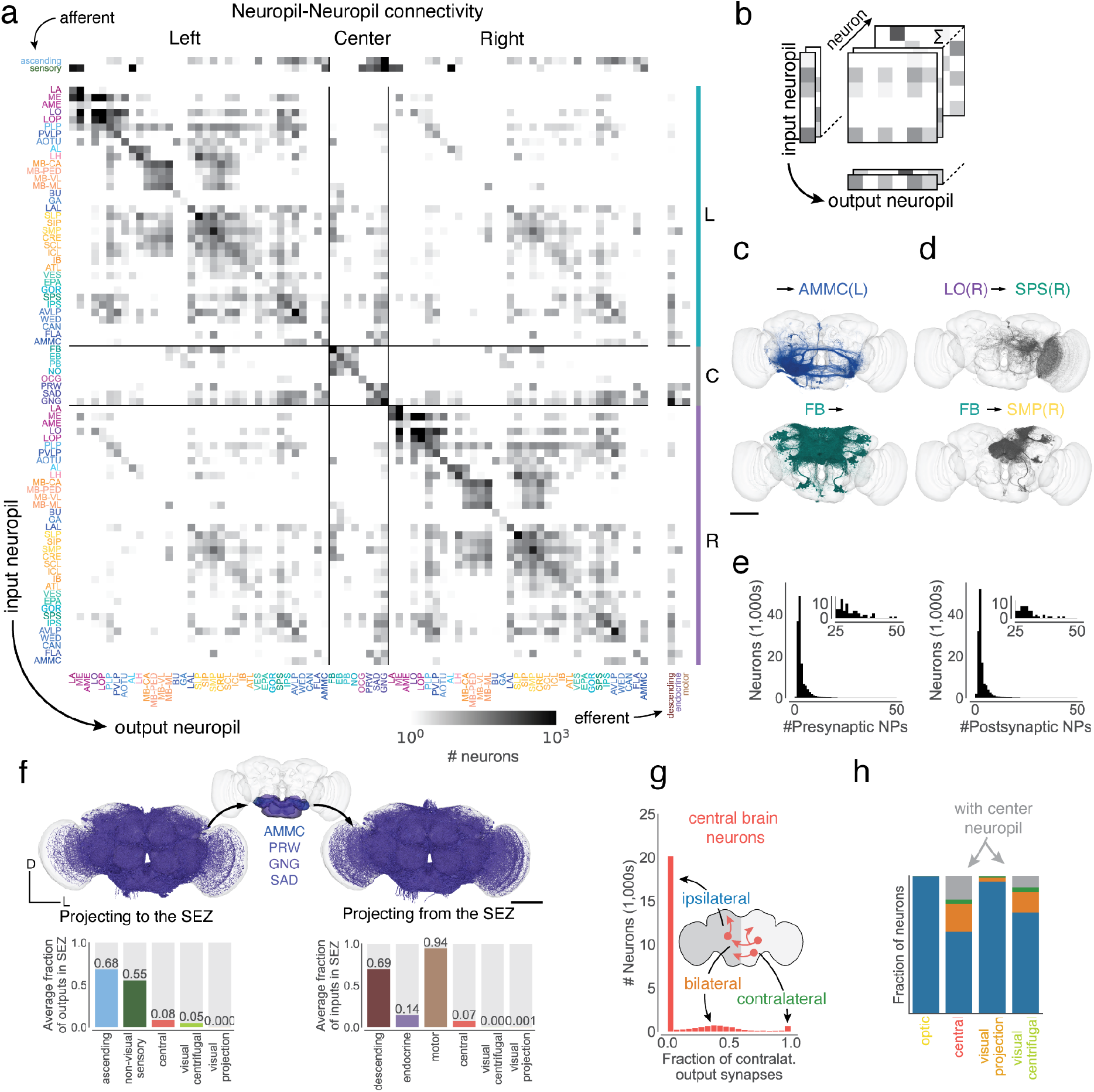
Neuropil projections and analysis of crossing neurons. (a) Whole brain neuropil-neuropil connectivity matrix. The main matrix was generated from intrinsic neurons, and afferent and efferent neuron classes are shown on the side. Incoming synapses onto afferent neurons and outgoing synapses from efferent neurons were not considered for this matrix. See Ext. Data Fig. 4-1 for neurotransmitter specific matrices. (b) Cartoon describing the generation of the matrix in (a) Each neuron’s connectivity is mapped onto synaptic projections between different neuropils. (c) shows examples from the matrix with each render corresponding to one row or column in the matrix and (d) shows examples from the matrix with each render corresponding to one square in the matrix. (e) Most neurons have pre- and postsynaptic locations in less than four neuropils. (f) Renderings (subset of 3,000 each) and input and output fractions of neurons projecting to (N=11916) and from (N=7528) the SEZ. The SEZ is roughly composed of five neuropils (the AMMC has a left and right homologue). Average input and output fractions were computed by summing the row and column values of the SEZ neuropils in the super-class specific projection matrices. (g) Fraction of contralateral synapses for each central brain neuron. (h) Fraction of ipsilateral, bilateral, contralateral, neurons projecting to and from the center neuropils per super-class. Scale bars: 100 µm

The projectome is a neuropil-neuropil matrix computed as follows. Each intrinsic neuron contributes to the projections between neuropils where it has pre- and postsynaptic sites. We weighted neuron projections by the product of the respective number of synapses and normalized the result for every neuron such that the matrix sums to the total number of intrinsic neurons. Each column corresponds to all the neurons projecting to a neuropil and each row to all neurons projecting out of it (Fig. 4b). Each square then represents the summed fractional weight of all neurons projecting between two neuropils (Fig. 4c,d). We added afferent and efferent neurons to the matrix by calculating the sum of the weighted neuron projections per super-class to and from all neuropils respectively.

While each neuropil is connected to many others, most neurons have synaptic sites in only a few neuropils (Fig. 4e). We repeated this process for each fast neurotransmitter type (Ext. Fig. 4-1). Some neuropil-neuropil connections exist strongly for one neurotransmitter but not others. For example, the neuropils making up the central complex (FP, EB, PB, NO) and the mushroom body (MB-CA, MB-PED, MB-VL, MB-ML) are largely tied together by excitatory connections.

We observed a strong symmetry between projections in the left and right hemisphere as well as with the central neuropils located on the midline (Ext. Data Fig. 4-2a,b); this highlights the strong similarity between the two sides of the brain. We observed that contralateral projections (projections from one side of the brain to the other) were generally weaker than projections to the same or ipsilateral neuropil (Ext. Data Fig. 4-2c).

The SEZ (Fig. 4f) is the ventral portion of the central brain, and has been shown to contribute to a variety of behaviors^69^. It is almost wholly unrepresented in the hemibrain reconstruction^33^, and is only partially reconstructed in the larval brain^39,111^. The five neuropils in the SEZ (left and right AMMC, GNG, SAD, and PRW; Fig. 4f) amount to 17.8% of central brain neuropil volume (0.0018 mm^3^ of 0.0103 mm^3^); they contain afferents mostly from non-visual sensory neurons (mechanosensory and taste) and ascending neurons, as well as a large number of efferents (motor, endocrine, and descending neurons - in fact, descending neurons receive on average 69% of their inputs in one of the five SEZ neuropils). The SEZ is thus important for information flow to and from the brain. Judging from the projectome (Fig. 4a), the SEZ neuropils interact with almost all parts of the brain. Notable exceptions are the central complex (EB, FB, PB, and NO) and the mushroom body (MB), suggesting less crosstalk between those circuits and neurons in the SEZ (explored in more detail in Fig. 6).

### Hemispheric organization

Our reconstruction includes both left and right brain hemispheres. This is important for tracing sensorimotor pathways that cross from one side to the other, and more generally for understanding interactions between the two hemispheres. The projectome (Fig. 4a) already reveals that most projections (88%) are ipsilateral or between neuropils on the same side of the brain.

The low fraction of non-ipsilateral neurons is primarily due to their scarceness in the optic lobes. Only 157 neurons (0.2%) in the optic lobes cross hemispheres, and cross the central brain without making synapses there (Supplemental Information 2) - these neurons are considered to be “fully contained” in the optic lobes because our definition depends only on synapse locations. These neurons mediate direct interactions between the two optic lobes, and their rarity suggests that these interactions represent a smaller fraction of the computations that occur within the optic lobes. Integration of information from both eyes may rely more on the abundant crossing connections between the central brain targets (AOTU, PLP, PVLP) of VPNs.

A higher proportion (40%) of central brain neurons are non-ipsilateral, largely owing to central neuropils, like those of the central complex and SEZ. To classify non-ipsilateral neurons, we started by examining the spatial distributions of their postsynapses (inputs). We divided the neuropils into three categories. Left and Right included the neuropils that come in mirror-symmetric pairs. Center included the seven remaining neuropils that are located on the midline. For each neuron, we computed the proportions of its postsynapses in Left, Right, and Center neuropils (Ext. Fig. 4-3). Each neuron was assigned to the dominant category, and near-ties were rare. The exceptions are symmetric neurons with cell bodies at the midline of the brain (Ext. Data Fig. 4-4, N=106).

Next, we asked how many neurons of Left and Right categories have presynapses (outputs) in the other hemisphere. Similar to the analysis of the 1st instar larval connectome^39^, we found that neurons projecting to the other hemisphere can be grouped into bilateral neurons, those with outputs in both hemispheres, and contralateral neurons which almost exclusively had presynapses in the other hemisphere (Fig. 4g-i). Notably, a much larger fraction of visual centrifugal neurons projected to the contralateral hemisphere than visual projection neurons, and both visual centrifugal neurons and neurons of the central brain contain a large fraction of bilateral neurons (Fig. 4h) - as stated earlier, this analysis again revealed the dominance of ipsilateral connections in the brain. While mixing between the hemispheres is more rare, mixing between sensory modalities within a hemisphere is common (see Fig. 6 below).

Many types of fly neurons are known to exhibit striking stereotypy across individuals, and also across both hemispheres of the same individual. A companion paper shows quantitatively using FlyWire and hemibrain data that these two kinds of stereotypy are similar in degree^44^.

### Optic lobes: columns and beyond

So far we have mentioned neurons that connect the optic lobes with each other, or with the central brain. The intricate circuitry within each optic lobe is also included in FlyWire’s connectome. Photoreceptor axons terminate in the lamina and medulla, neuropils of the optic lobes (Fig. 5a,b). Each eye contains approximately 800 ommatidia that map to columns in the lamina arranged in a hexagonal lattice (Fig. 5b). This structure repeats in subsequent neuropils from lamina to medulla to lobula to lobula plate. The neuropils have been finely subdivided into layers that are perpendicular to the columns^112^. The 2D visual field is mapped onto each layer. Any given cell type tends to synapse in some subset of the layers. Cell types vary greatly in size. Uni-columnar cell types are the smallest (Fig. 5b,c). At the other extreme are large cells that span almost all columns (Fig. 5d). In between there are many multi-columnar cell types that are still being classified (Fig. 5e).

**Figure 5:**
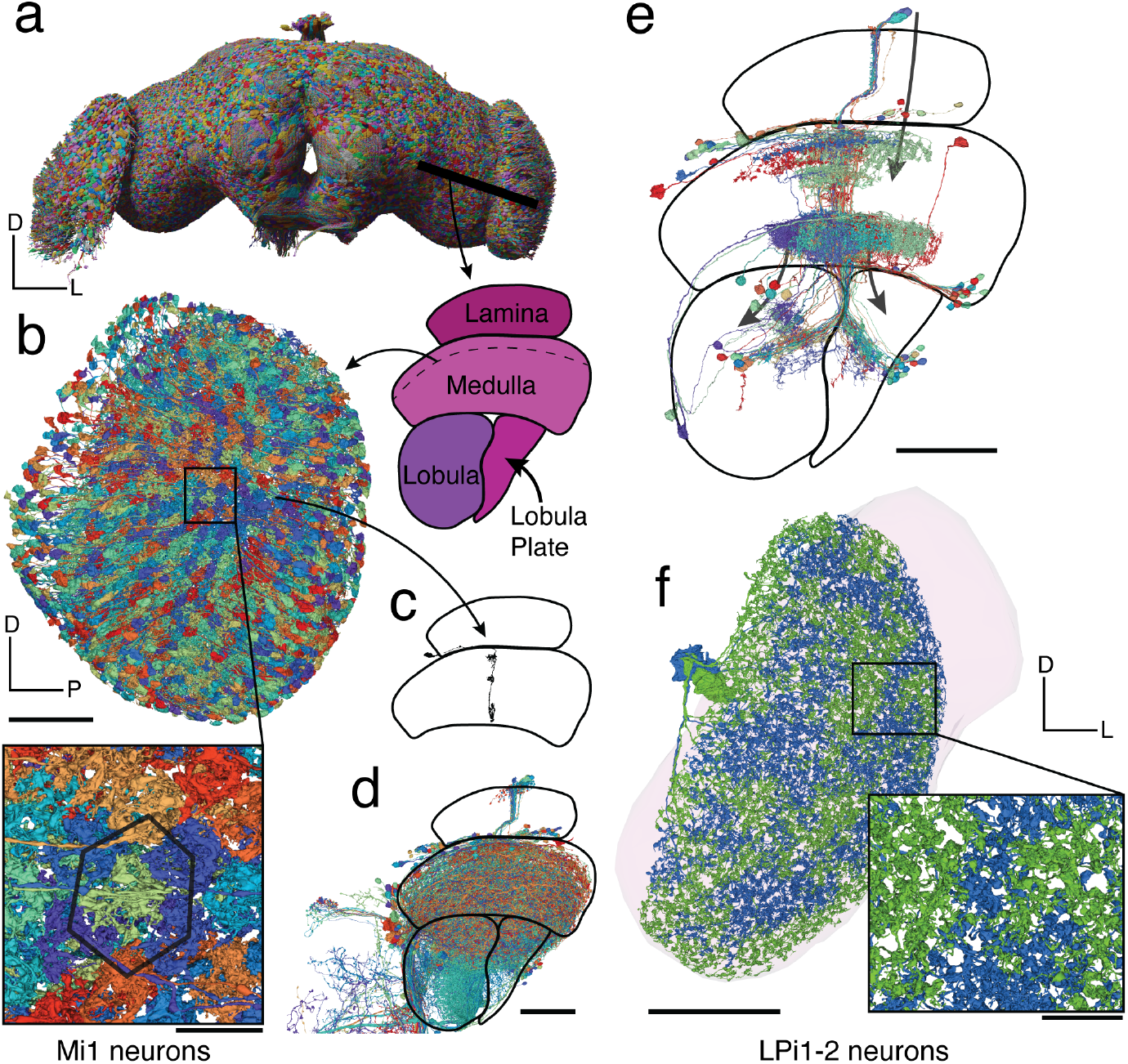
Optic lobes. (a) Rendering of a subset of the neurons in the fly brain. A cut through the optic lobe is highlighted. (b) All 779 Mi1 neurons in the right optic lobe. (c) A single Mi1 neuron, (d) all neurons crossing through the column in c as defined by a cylinder in the medulla with 1 µm radius through it, and (e) all neurons sharing a connection with the single Mi1 neuron shown in (c) (≥ 5 synapses) - 3 large neurons (CT1, OA-AL2b2, Dm17) were excluded for the visualization. (f) The two LPi1-2 neurons in the right lobula plate (neuropil shown in background). Scale bars: 50 µm (b,c,d,e,f), 10 µm (b-inset)

Mi1 is a true “tiling” type, i.e., its arbors cover the visual field with little or no overlap, and have similar size and shape (Fig. 5b). Dm12 arbors overlap with each other, but the spatial arrangement is still regular. These and other distal medullary cell types were previously characterized by multicolor light microscopy^113^. Our EM reconstructions reveal even more detailed information about the spatial patterning of these types (e.g., co-fasciculation of neurites of neighboring Dm12 cells). More importantly, FlyWire’s reconstruction encompasses all multi-columnar cell types, including those outside the medulla. Judging from the many examples we have studied throughout the optic lobe, it seems that regular coverage of the visual field without gaps is a defining criterion for most cell types, similar to mammalian retina^114^. There are, however, exceptional cell types that cover the visual field in an irregular manner. For example, there are exactly two LPi1-2 cells per optic lobe^43^. The shapes of each pair are complementary, as if they were created by cutting the visual field into two pieces with a jigsaw (Fig. 5f); this tiling was not evident when reconstructing only a portion of an optic lobe^43^.

Much of the existing research on widefield visual motion processing has relied on the simplifying idea that the computations are mostly in columnar circuits, and the columnar outputs are finally integrated by large tangential cells in the lobula plate. This research has been aided by wiring diagrams containing connections between cells in the same column or neighboring columns^16,17,42^. An absence of information across columns, has necessitated treating each column as identical in simulations of the optic lobe^115^. FlyWire’s connectome contains not only the columnar neurons (Fig. 5b), but also all neurons that extend across columns (Fig. 5d,e). These neurons are both excitatory and inhibitory, and can support interactions between even distant columns. This opens up the possibility of a much richer understanding of optic lobe computations, and this is explored in a companion paper on hue selectivity (Christenson et al. *in prep*).

Some columnar cell types are known to exhibit spatial gradients in connectivity ^116^, and our reconstruction makes it possible to investigate such gradients for any columnar cell type in the optic lobe. Similar gradients have also been studied in mammalian retina ^117^, and such continuous variation is an interesting complement to the conventional notion that cell types are discrete.

### Analysis of information flow

While afferent and efferent neurons make up a numerically small proportion of the brain (estimated 14.7% and 1.1% respectively), they are important because they connect the brain to the outside world. Examining connections to these neurons is useful when attempting to predict the functions of intrinsic neurons from the connectome. For example, one might try to identify the shortest path in the connectome from an afferent (input) neuron that leads to a given intrinsic neuron. The sensory modality of the afferent neuron could provide a clue as to the function of the intrinsic neuron. This approach, while intuitive, ignores connection strengths and multiplicities of parallel pathways. We therefore use a probabilistic model^20^ to estimate information flow in the connectome, starting from a set of seed neurons (Fig. 6a; see Methods).

**Figure 6.**
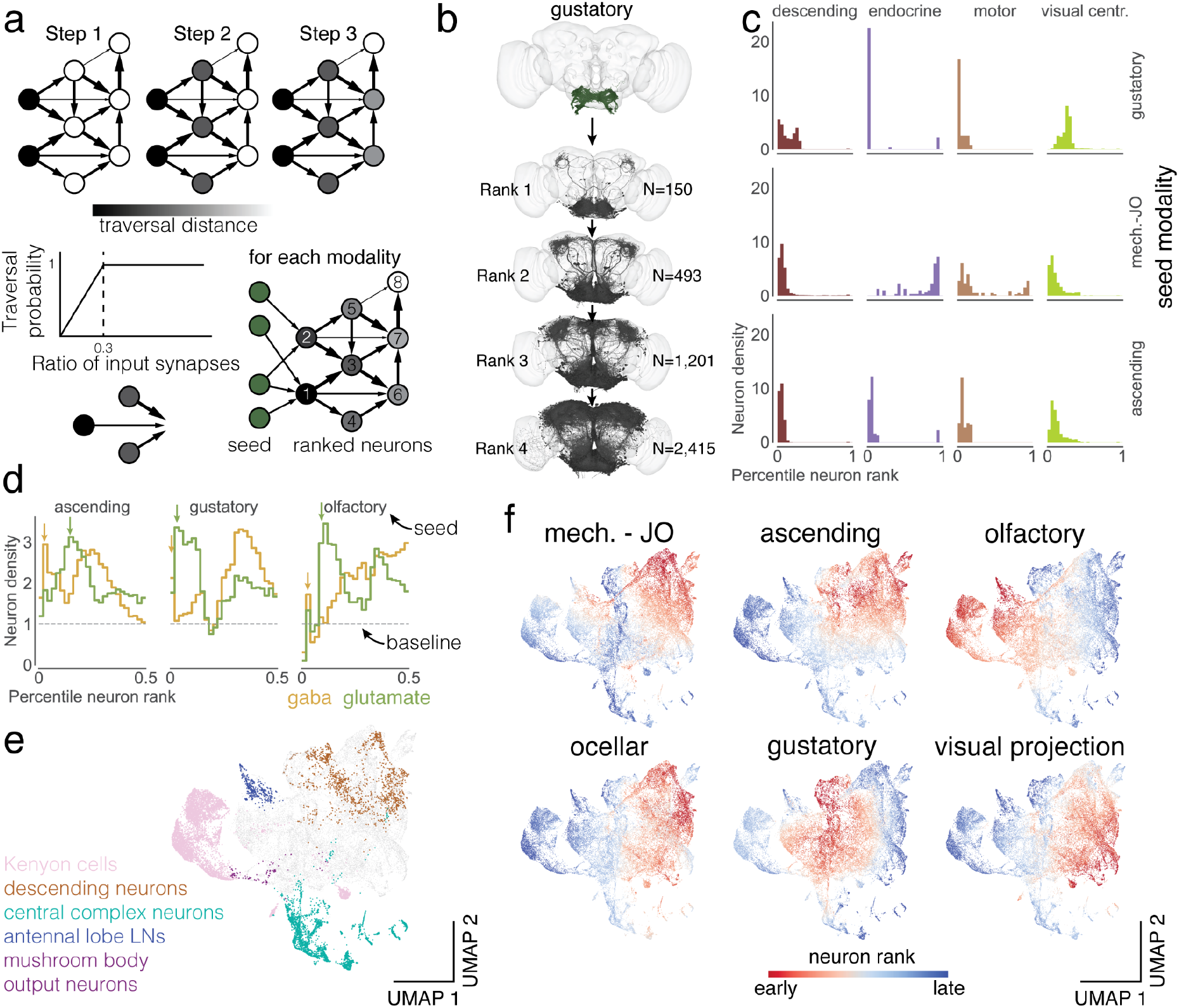
Information flow through the *Drosophila* central brain. (a) We applied the information flow model for connectomes by Schlegel et al. ^20^ to the connectome of the central brain neurons. Neurons are traversed probabilistically according to the ratio of incoming synapses from neurons that are in the traversed set. The information flow calculations were seeded with the afferent classes of neurons (including the sensory categories). (b) We rounded the traversal distances to assign neurons to layers. For gustatory neurons, we show a subset of the neurons (up to 1,000) that are reached in each layer. (c) For each sensory modality we used the traversal distances to establish a neuron ranking. Each panel shows the distributions of neurons of each super-class within the sensory modality specific rankings (see Ext. Data Fig. 6-1a for the complete set). (d) We assign neurons to neurotransmitter types and show their distribution within the traversal rankings similar to (c). The arrows highlight the sequence of GABA - glutamate peaks found for almost all sensory modalities (see Ext. Data Fig. 6-1b for the complete set). (e) We UMAP projected the matrix of traversal distances to obtain a 2d representation of each neuron in the central brain. Neurons from the same class co-locate (see also Ext. Data Fig. 6-2) (f) Neurons in the UMAP plot are colored by the rank order in which they are reached from a given seed neuron set. Red neurons are reached earlier than blue neurons (see Ext. Data Fig. 6-1c for the complete set).

The likelihood of a neuron being traversed increases with the fraction of inputs from already traversed neurons and caps out at an input fraction of 30%. We ran the traversal model for every subset of afferent neurons as seeds (N=12 input modalities to the central brain, Fig. 6b, Fig. 2e, Supplemental Information 3, see Methods for full list). We then measured information flow from these starting neurons to all intrinsic and efferent neurons of the central brain (for this analysis, we ignore circuitry within the optic lobes, and consider VCNs (visual centrifugal neurons) as efferents of the central brain). We then ranked all neurons by their traversal distance from each set of starting neurons and normalized the order to percentiles. For instance, a neuron at the 20th percentile had a lower rank than 80% of neurons. This allowed us to determine how early information from each afferent modality reached various targets, including the descending neurons, endocrine neurons, motor neurons and visual centrifugal neurons (Fig. 6c, Ext. Data Fig. 6-1a). As expected, endocrine neurons are closest to the gustatory sensory neurons while motor and descending neurons were reached early for mechanosensory and visual afferents (Ext. Data Fig. 6-1a).

Do the afferent cell classes target inhibitory neurons early or late? We found that putative inhibitory neurons (neurons predicted to express GABA and glutamate) were overrepresented in the set of early neurons (Fig. 6d). Surprisingly, we identified a sequence of GABAergic and glutamatergic peaks in the sequence of neurons targeted that was replicated for almost all afferent modalities (Ext. Data Fig. 6-1b).

To visualize information flow in a common space, we treated the traversal distances starting from each seed population as a neuron embedding and built a UMAP projection from all of these embeddings (Fig. 6e). Within the map, we found that neurons of the same cell class (e.g. two groups of Kenyon cells, all mushroom body output neurons, all antennal lobe local neurons, and all central complex neurons) are clustered. Next, we displayed traversal order on top of the UMAP plot to compare traversal orders starting from different modalities. We find that every neuron in the central brain can be reached by starting from any modality - this “small world” property of the network is covered in more detail in a companion paper (Lin et al., *in prep*). Comparing orders revealed that almost all neurons in the central brain are reached early starting from some modality, with the exception of neurons in the central complex (Fig. 6f, Ext. Data Fig. 6-2), highlighting that the central complex is dominated by internal computations^15^. Kenyon cells were contained in two clusters - one of which is targeted very early from olfactory receptor neurons and the other targeted early by visual projection neurons^118^.

Our information flow analysis provides a compressed representation of the connectome, but ignores signs of connections and the biophysics of neurons and synapses, and therefore terms like “early” and “late” should not be interpreted as true latencies to sensory stimulation. A companion paper ^40^ builds a leaky integrate-and-fire model of *Drosophila* brain dynamics, using the connectome and including connection weights (number of synapses) and putative connection signs (excitatory or inhibitory).

### Cell types and other annotations

Neurons in *Drosophila* are considered to be identifiable across hemispheres and individuals^119,120^, enabling cell type classification of all neurons in FlyWire. Such classification is useful for generating testable hypotheses about circuit function from the connectome. FlyWire community members, many experts in diverse regions of the fly brain, have shared 91,649 annotations of 59,548 neurons (Supplemental Information 4), including the majority of sexually-dimorphic neurons (Deutsch et al., *in prep*), sensory neurons^41^, as well as a diversity of cell types in the optic lobes and SEZ (Fig. 2f). Each neuron in FlyWire is also given a unique identifier based on the neuropil it receives and sends most of its information. Curation of these annotations continues, and we invite further community efforts to identify cell types, which can be contributed through Codex (codex.flywire.ai).

In addition, matching between cell types identified in the hemibrain^33^ and both hemispheres of FlyWire provides additional annotations for neurons contained in both datasets. Our companion paper^44^ provides cell type annotations for 26,150 neurons via such matching. However, many types proposed in the hemibrain reconstruction could not yet be re-identified in the FlyWire dataset.

All cell annotations can be queried in Codex. Some of these have already been mentioned, such as the “flow” annotations of intrinsic vs. afferent vs. efferent, super-class annotations of Fig. 2, neurotransmitter predictions, left-right annotations for cell body location, in addition to lineages, or groups of neurons derived from a single neuroblast^121^.

### Ocellar circuit structure and function: linking sensory inputs to motor outputs

The completeness of the FlyWire connectome enables tracing complete pathways from sensory inputs to motor outputs. Here we demonstrate this capability by examining circuits that emanate from the ocellar ganglion and leveraging cell type information. In addition to the large compound eyes, flying insects have smaller visual sensory organs^122^, including the three ocelli on the dorsal surface of the head cuticle (Fig. 7a). The ocelli are under-focused eyes, projecting a blurry image of light level changes in the UV and blue color spectrum^123,124^; these eyes are thought to be useful for flight control and orientation relative to the horizon ^125^. Importantly, while the role of the ocelli has been hypothesized (e.g., light level differences between the eyes when the fly is shifted off axis should quickly drive righting motions of the head, wings, and body to stabilize gaze and re-orient the body), little is known about the circuitry downstream of this sensory organ that would mediate this function.

**Figure 7.**
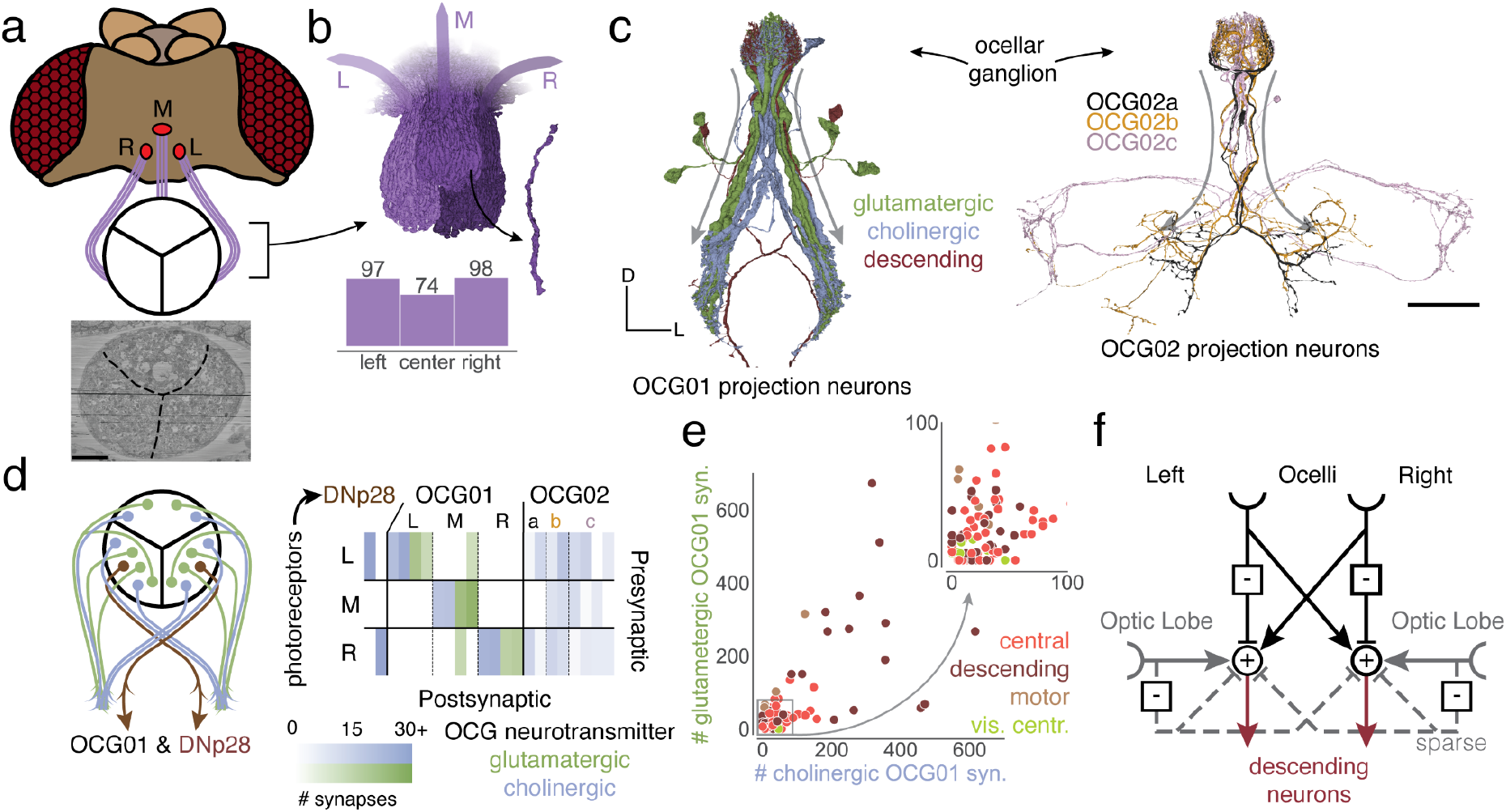
Ocellar circuits and their integration with visual projection neurons. (a) Overview of the three ocelli (left, medial, right) which are positioned on the top of the head. Photoreceptors from each ocellus project to a specific subregion of the ocellar ganglion which are separated by glia (marked with black lines on the EM). (b) Renderings of the axons of the photoreceptors and their counts, and (c) OCG01, OCG02 and DNp28 neurons with arbors. “Information flow” from pre- and postsynapses is indicated by arrows along the arbors. (d) Connectivity matrix of connections between photoreceptors and ocellar projection neurons, including two descending neurons (DNp28). (e) Comparison of number of glutamatergic and cholinergic synapses from ocellar projection neurons onto downstream neurons colored by super-class (R=.65, p<1e-21). (f) Summary of the observed connectivity between ocellar projection neurons, visual projection neurons and descending neurons. Scale bar: 100 µm

Photoreceptor axons (N=270) from the three ocelli innervate three distinct regions of the ocellar ganglion separated by glial sheets (Fig. 7a, b). The ocellar ganglion additionally contains 62 neurons that we categorized into four broad groups (Fig. 7c, Ext. Data Fig. 7-1a): local neurons (N=15), two types of interneurons, divided based on their arborizations and caliber (OCG01 (N=12), OCG02 (N=8)), descending neurons (DNp28, N=2), and centrifugal or feedback neurons (N=25). Ocellar local neurons are small (116 outgoing synapses, 449 µm path length on average) and connect sparsely with photoreceptors from all ocelli.

Twelve OCG01 interneurons and two descending neurons (DNp28, one per lateral ocellus) represent the main pathway from the ocellar ganglion to the central brain. DNp28 projects to the intermediate, haltere, wing, and neck tectula of the ventral nerve cord^62,71^. In each ocellus, half of the OCG01s were inferred to express glutamate (likely inhibitory), and the other half acetylcholine (excitatory). There are four OCG01s per ocellus (Fig. 7d). OCG01s tile the ocellar ganglion, indicating their receptive fields tile the visual fields of the ocelli (Ext. Data Fig. 7-1 b,c). OCG02 axons are much thinner than the OCG01s, and likely transmit signals slower^126^. Two OCG02 subgroups (a, b) innervate similar neuropils to the OCG01s (IPS, SPS), and OCG02c neurons target the PLP, a brain region that also receives input from visual projection neurons from the compound eyes^72^.

Neurons downstream from OCG01s in the IPS, SPS, and GNG receive inhibitory input from the ipsilateral ocellus and excitatory input from the contralateral ocellus (Fig. 7d, right), and the amount of synaptic input from each ocellus is tightly correlated (Fig. 7e, R=0.65, p<1e-21) - this balance is likely to be a key ingredient in how signals are integrated (the descending circuits are activated by a signal difference between the eyes). We found that 15 different descending neurons (DNs) each receive over 200 synapses from the OCG01 neurons. For example, two DNs in each hemisphere received over 30% of their synaptic inputs in the brain from ocellar projection neurons: DNp20/DNOVS1 (left: 57%, right: 44%), DNp22/DNOVS2 (left: 36%, right: 33%). DNOVS1 and other descending neurons with strong input from OCG01s generally receive strong input from ipsilateral visual projection neurons as well (Ext. Data Fig. 7-1d). For example, DNOVS1 is also activated by rotational optic flow fields across the compound eye, and projects to the neck motor system^127,128^. A handful of glutamatergic (putative inhibitory) visual projection neurons sparsely innervate descending neurons in both hemispheres. As the ocelli transmit mainly information about light levels, the dense integration with motion direction signals from the compound eyes was not previously appreciated, but should aid in precision adjustments of head and body movements for gaze stabilization and flight control^129^.

There is also extensive feedback from the brain directly to the ocellar ganglion via 25 ocellar centrifugal neurons (OCC). We found striking targeting specificity of two OCC subgroups (OCC01a, b) which synapse onto all OCG01 and DNp28 neurons with strong connections compared with their overall synaptic budget (Ext. Data Fig. 7-1e). The OCC01s receive input in a wide range of neuropils, notably the SEZ, as well as IPS and SPS, the same neuropils that receive inputs from the OCG projection neurons (Ext. Data Fig. 7-1f). It remains to be determined what role the OCCs play in gating visual information and potentially driving the OCGs in the absence of photoreceptor activity.

Based on the summary wiring diagram of Fig. 7f, we hypothesize how the pathways from the ocelli to descending neurons function. As in a Braitenberg vehicle for phototaxis^130^, excitation and inhibition are organized so that the head and body of the fly should roll around the anteroposterior axis to orient the ocelli towards light. In this example, the whole-brain connectome, extending from brain inputs to outputs, uncovers new pathways and facilitates the generation of testable hypotheses for circuit mechanisms of sensorimotor behavior.

## Discussion

By reconstructing a complete brain wiring diagram, FlyWire enables many kinds of studies that were not previously possible using wiring diagrams of portions of the fly brain. The optic lobes and the SEZ are two prominent regions mostly missing from the hemibrain, the previous state of the art. Both sides of the brain are included, enabling the tracing of pathways that cross the midline. Due to the presence of afferent and efferent neurons, one can trace pathways from sensory inputs to intrinsic neurons to brain outputs (motor, endocrine, and descending neurons). This was done in a global fashion using the information flow model, and more specifically to uncover the structure and hypothesize a circuit mechanism for behaviors supported by the ocelli. Our companion papers provide additional global analyses of the connectome (Lin et al., *in prep*) and studies of specific families of pathways.

### Connectome annotation

Connectome annotation with structural and functional information is an important emerging field, analogous to genome annotation. Annotations are important because they make the connectome usable for hypothesis generation about circuit function. We carried out a hierarchical and systematic annotation of all neurons in the connectome as detailed in our companion paper^44^, describing over 4000 robustly identifiable cell types. We also collected a large number of annotations from the community (57% of all neurons have an annotation label) leveraging a broad knowledge base. Further curation of these labels will help to refine them.

### Comparative connectomics

For the first time, one can now compare entire connectomes of different species, starting with *Drosophila melanogaster* and *C. elegans*, as touched on by the present manuscript, and explored in more depth by Lin et al (Lin et al., *in prep*). One can also compare connectomes of the same species at different developmental stages^39^. While ours is still the only adult fly connectome, it can be compared with the hemibrain reconstruction where they overlap, to detect wiring differences between adults of the same species, and to validate and extend cell type definitions^44^.

### Connectomes, transcriptomes, and brain development

Transcriptomics with single cell resolution is being applied to mammalian brains^131^, and to the *Drosophila* brain as well. Transcriptomic atlases of the central adult brain^95,132^ and optic lobes^133,134^ are appearing. Comparing connectomes with transcriptomes is already proving useful for studying molecular mechanisms of development^135–137^. Clearly more fly connectomes at multiple developmental stages are needed.

### Brain simulation

Connectome-based brain simulation was one of the original motivations for connectomics^138^. A neural network simulation of visual motion detection based on the wiring diagram of columnar circuits in the optic lobe has been created^115^. Such a connectome-based approach can at last be scaled up to an entire brain^40,139^.

### Block face versus serial section EM

The hemibrain was reconstructed^33^ from images acquired by FIB-SEM^140–142^, a form of block face EM^143,144^. FlyWire is based on transmission EM images of serial sections (ssTEM) that were manually cut and collected, and then automatically imaged^52^, an evolution of the approach that was used for the *C. elegans* connectome^36^. In the end, both block face and serial section EM have turned out to be viable for fly connectomes. Both approaches yield similar accuracy (Ext. Fig. 1-2, Ext. Fig. 1-3). Hybrid methods that combine both imaging approaches are also being developed^142^.

### Artificial and human intelligence

Owing to the use of artificial intelligence (AI), the hemibrain and FlyWire have yielded connectomes that are orders of magnitude larger than those of *C. elegans*^38^ or the larval fly^39^. The hemibrain images were automatically segmented using flood-filling convolutional nets^145^, whereas FlyWire used the older approach of boundary-detecting convolutional nets^146,147^. FlyWire also required another kind of AI, alignment of serial section images using convolutional nets^53^. While the hemibrain used custom software to achieve 3D alignment of volume EM data^148,149^, AI-based alignment was crucial for making ssTEM as amenable to automated reconstruction^150^. In spite of enormous progress in AI, both the hemibrain^33^ and FlyWire (Methods) required an estimated 50 and 30 person-years of human effort for proofreading the automated segmentation respectively (see Methods). This is because AI has reduced the amount of human labor required per unit brain volume, but EM image volumes have increased even faster. Further reduction in human proofreading is necessary for reconstructing many fly connectomes to study variation, or to scale up to whole mammalian brains.

### Imaging smaller

The EM images used by FlyWire were acquired at a resolution of 4×4×40 nm^3^. Increasing this resolution would presumably enable accurate attachment of twigs to backbones, which is currently the main factor limiting the accuracy of reconstructing synaptic connectivity. Higher resolution might also enable the reconstruction of electrical synapses, which are included in the *C. elegans* connectome. However, the lower limit of the size of functional electrical synapses is unknown^151^, raising the possibility that no current volume EM method can capture all electrical synapse connectivity. Increasing resolution by 2× in all three dimensions would increase the data volume by 8×. Handling much larger data volumes should be possible as methods for acquiring and analyzing EM images are progressing rapidly.

### Imaging larger

Imaging a larger volume would open up other interesting opportunities. Imaging a whole fly CNS would enable the mapping of all pathways linking the brain and VNC. In the meantime, it is possible to establish correspondences between FlyWire and FANC, a reconstruction of a separate VNC^61,70^. The first *C. elegans* connectome was obtained similarly as a mosaic drawn from multiple worms ^36^. Imaging an entire fly, both CNS and body, would enable the addition of sensory organs and muscles to the reconstruction. This also has precedent in the *C. elegans* connectome^38^, which includes neuromuscular junctions, the *Platynereis dumerilii* larva^152^, and the 1st instar *Drosophila* larva for which a whole-animal EM dataset was recently published^153^.

FlyWire and other related technologies have already been applied to millimeter-scale chunks of mammalian brain^24,25^, which are >50× larger in volume than a fly brain. The U.S. National Institutes of Health is planning a ten year project to reconstruct a whole mouse brain from an exabyte of EM images and a report from the Wellcome trust recently examined the road to a whole mouse brain connectome^154^.

### Openness

The 1996 Bermuda Principles mandated daily release of Human Genome Project sequences into the public domain^155^. We believe that openness is also important for large-scale connectomics projects, particularly because these projects are expensive, require coordinated effort, and take several years to complete - sharing connectomes only after proofreading and annotation are completed prevents scientific discovery that can occur while the connectome is being completed. Shortly after its inception, FlyWire has been open to any *Drosophila* researcher. As a result, hundreds of scientists and proofreaders from over 50 labs joined FlyWire with over 200 of them contributing over 100 edits (Supplemental Table 1) and 86 contributing ten or more annotations (Supplemental Table 2). As a result, there are multiple studies that used completed portions of FlyWire’s connectome as proofreading proceeded^13,18,20,41,51,69,156–163^. Openness has also enabled FlyWire to move faster by incorporating data sources from the community. The EM data on which FlyWire is built was shared in 2018 by Bock and colleagues^52^. FlyWire’s synapse data was previously published by Buhmann et al.^45^ who incorporated synapse segmentations from Heinrich et al.^46^, neurotransmitter labels for every synapse were made available ahead of publication by Eckstein et al.^47^, numerous annotations were contributed by Schlegel et al., and over 90K (and counting) cell annotations have been shared by the community. Overall we anticipate that similar approaches based on an open ecosystem will allow connectomics to scale more efficiently, economically, and equitably.

## FlyWire Consortium

Doug Bland^1^, Krzysztof Kruk^3^, Zairene Lenizo^16^, Alexander Shakeel Bates^4,5,12,13^, Nseraf^3^, Austin T. Burke^1^, Katharina Eichler^5^, Nashra Hadjerol^16^, Kyle Patrick Willie^1^, Ryan Willie^1^, Yijie Yin^5^, John Anthony Ocho^16^, Sven Dorkenwald^1,2^, Joshua Bañez^16^, Arti Yadav^17^, Shirleyjoy Serona^16^, Rey Adrian Candilada^16^, Dustin Garner^18^, Philipp Schlegel^4,5^, Jet Ivan Dolorosa^16^, Ariel Dagohoy^16^, Remer Tancontian^16^, Mendell Lopez^16^, Regine Salem^16^, Griffin Badalamente^5^, annkri (Anne Kristiansen)^3^, Kendrick Joules Vinson^16^, Nelsie Panes^16^, Laia Serratosa Capdevila^5^, Anjali Pandey^17^, Darrel Jay Akiatan^16^, Ben Silverman^1^, Dharini Sapkal^17^, Shaina Mae Monungolh^16^, Jay Gager^1^, Varun Sane^5^, Miguel Albero^16^, AzureJay (Jaime Skelton)^3^, Márcia dos Santos^5^, David Deutsch^1,9^, Zeba Vohra^17^, Kaiyu Wang^14^, Emil Kind^19^, Chitra Nair^17^, Dhwani Patel^17^, Imaan F. M. Tamimi^5^, Michelle Darapan Pantujan^16^, James Hebditch^1^, Alexandre Javier^5^, Rashmita Rana^17^, Bhargavi Parmar^17^, Merlin Moore^1^, Mark Lloyd Pielago^16^, Allien Mae Gogo^16^, Markus William Pleijzier^4^, Mark Larson^20^, Joseph Hsu^5^, Thomas Stocks^3^, Jacquilyn Laude^16^, Itisha Joshi^17^, Chereb Martinez^16^, Dhara Kakadiya^17^, John David Asis^16^, Amalia Braun^21^, Clyde Angelo Lim^16^, Alvin Josh Mandahay^16^, Marchan Manaytay^16^, Marina Gkantia^5^, Kaushik Parmar^17^, Quinn Vanderbeck^12^, Claire E. McKellar^1^, Philip Lenard Ampo^16^, Daril Bautista^16^, Irene Salgarella^5^, Christopher Dunne^5^, John Clyde Saguimpa^16^, Eva Munnelly^5^, Chan Hyuk Kang^22^, Jansen Seguido^16^, Jinmook Kim^22^, Gizem Sancer^23^, Lucia Kmecova^24^, Christa Baker^1^, Jenna Joroff^12^, Steven Calle^24^, Cathy Pilapil^16^, Yashvi Patel^17^, Olivia Sato^20^, Siqi Fang^5^, Paul Brooks^5^, Mai Bui^25^, JousterL (Matthew Lichtenberger)^3^, edmark tamboboy^16^, Katie Molloy^20^, Alexis E Santana-Cruz^24^, Janice Salocot^16^, Celia David^1^, Kfay^3^, Seongbong Yu^22^, Arzoo Diwan^17^, Farzaan Salman^26^, Szi-chieh Yu^1^, Monika Patel^17^, TR77^3^, Sarah Morejohn^1^, Sebastian Molina-Obando^27^, Sanna Koskela^14^, Tansy Yang^14^, bl4ckscor3 (Daniel Lehmann)^3^, Sangeeta Sisodiya^17^, Selden Koolman^1^, Philip K. Shiu^28^, Sky Cho^25^, Brian Reicher^20^, Marlon Blanquart^5^, Marissa Sorek^1,3^, Lucy Houghton^18^, Hyungjun Choi^22^, Matt Collie^20^, Joanna Eckhardt^1^, Benjamin Gorko^18^, Li Guo^18^, Zhihao Zheng^1^, Alisa Poh^29^, Marina Lin^25^, István Taisz^4^, Wes Murfin^52^, Álvaro Sanz Díez^37^, Peter Gibb^12^, Nils Reinhard^30^, Nidhi Patel^17^, Sandeep Kumar^1^, Minsik Yun^31^, Megan Wang^1^, Devon Jones^1^, Lucas Encarnacion-Rivera^32^, Annalena Oswald^27^, Akanksha Jadia^17^, Leonie Walter^19^, Nik Drummond5, Ibrahim Tastekin^33^, Xin Zhong^19^, Yuta Mabuchi^34^, Fernando J Figueroa Santiago^24^, Urja Verma^17^, Nick Byrne^20^, Edda Kunze^19^, Thomas Crahan^18^, Hewhoamareismyself (Ryan Margossian)^3^, Haein Kim^34^, Iliyan Georgiev^3^, Fabianna Szorenyi^24^, Benjamin Bargeron^35^, Tomke Stuerner^4,5^, Damian Demarest^36^, Atsuko Adachi^37^, Burak Gür^27^, Andrearwen^3^, Robert Turnbull^5^, a5hm0r^3^, Andrea Sandoval^28^, Diego A. Pacheco^12^, Haley Croke^38^, Alexander Thomson^14^, Jonas Chojetzki^27^, Connor Laughland^14^, Suchetana B. Dutta^19^, Paula Guiomar Alarcón de Antón^19^, Binglin Huang^18^, Patricia Pujols^24^, Isabel Haber^20^, Amanda González-Segarra^28^, Albert Lin^1,6^, Daniel T. Choe^39^, Veronika Lukyanova^40^, Marta Costa^5^, Maria Ioannidou^27^, Zequan Liu^41^, Tatsuo Okubo^12^, Miriam A. Flynn^14^, Gianna Vitelli^35^, Meghan Laturney^28^, Feng Li^14^, Shuo Cao^42^, Carolina Manyari-Diaz^35^, Hyunsoo Yim^22^, Anh Duc Le^38^, Kate Maier^35^, Seungyun Yu^22^, Yeonju Nam^22^, Mavil^3^, Nino Mancini^35^, Eleni Samara^21^, Amanda Abusaif^28^, Audrey Francis^43^, Jesse Gayk^17^, Sommer S. Huntress^44^, Raquel Barajas^33^, Mindy Kim^20^, Xinyue Cui^34^, Amy R Sterling^1,3^, Anna Li^12^, Gabriella R. Sterne^28^, Lena Lörsch^27^, Keehyun Park^22^, Alan Mathew^5^, 김진성^22^, Taewan Kim^22^, Guan-ting Wu^45^, Serene Dhawan^46^, Margarida Brotas^33^, Cheng-hao Zhang^45^, Shanice Bailey^5^, Alexander Del Toro^28^, Arie Matsliah^1^, Kisuk Lee^1,10^, Thomas Macrina^1,2^, Casey Schneider-Mizell^7^, Mert Erginkaya^33^, Sergiy Popovych^1,2^, Oluwaseun Ogedengbe^1^, Runzhe Yang^1,2^, Akhilesh Halageri^1^, Will Silversmith^1^, Stephan Gerhard^47^, Andrew Champion^4,5^, Nils Eckstein^14^, Dodam Ih^1^, Nico Kemnitz^1^, Manuel Castro^1^, Zhen Jia^1^, Jingpeng Wu^1^, Eric Mitchell^1^, Barak Nehoran^1,2^, Shang Mu^1^, J. Alexander Bae^1,11^, Ran Lu^1^, Eric Perlman^8^, Ryan Morey^1^, Kai Kuehner^1^, Derrick Brittain^7^, Chris S. Jordan^1^, David J. Anderson^42^, Rudy Behnia^37^, Salil S. Bidaye^35^, Davi D. Bock^15^, Alexander Borst^21^, Eugenia Chiappe^33^, Forrest Collman^7^, Kenneth J. Colodner^44^, Andrew Dacks^26^, Barry Dickson^14^, Jan Funke^14^, Denise Garcia^38^, Stefanie Hampel^24^, Volker Hartenstein^48^, Bassem Hassan^19^, Charlotte Helfrich-Forster^30^, Wolf Huetteroth^49^, Gregory S.X.E. Jefferis^4,5^, Jinseop Kim^22^, Sung Soo Kim^18^, Young-Joon Kim^31^, Wei-Chung Lee^12^, Gerit A. Linneweber^19^, Gaby Maimon^43^, Richard Mann^37^, Mala Murthy^1^, Michael Pankratz^36^, Lucia Prieto-Godino^46^, Jenny Read^40^, Michael Reiser^14^, Katie von Reyn^38^, Carlos Ribeiro^33^, Kristin Scott^28^, Andrew M. Seeds^24^, Mareike Selcho^49^, H. Sebastian Seung^1,2^, Marion Silies^27^, Julie Simpson^18^, Mathias F. Wernet^19^, Rachel I. Wilson^12^, Fred W. Wolf^50^, Zepeng Yao^51^, Nilay Yapici^34^, Meet Zandawala^30^

^16^SixEleven, Davao City, Philippines

^17^ariadne.ai ag, Buchrain, Switzerland

^18^University of California, Santa Barbara, USA

^19^Freie Universität Berlin, Berlin, Germany

^20^Harvard, Boston, USA

^21^Department Circuits-Computation-Models, Max Planck Institute for Biological Intelligence, Planegg, Germany

^22^Sungkyunkwan University, Seoul, South Korea

^23^Department of Neuroscience, Yale University, New Haven, USA

^24^Institute of Neurobiology, University of Puerto Rico Medical Sciences Campus, San Juan, Puerto Rico

^25^Program in Neuroscience and Behavior, Mount Holyoke College, South Hadley, USA

^26^Department of Biology, West Virginia University, Morgantown, USA

^27^Johannes-Gutenberg University Mainz, Mainz, Germany

^28^University of California, Berkeley, USA

^29^University of Queensland, Brisbane, Australia

^30^Julius-Maximilians-Universität Würzburg, Würzburg, Germany

^31^Gwangju Institute of Science and Technology, Gwangju, South Korea

^32^Stanford University School of Medicine, Stanford, USA

^33^Champalimaud Foundation, Lisbon, Portugal

^34^Cornell University, Ithaca, USA

^35^Max Planck Florida Institute for Neuroscience, Jupiter, USA

^36^University of Bonn, Bonn, Germany

^37^Zuckerman Institute, Columbia University, New York, USA

^38^Drexel, Philadelphia, USA

^39^Seoul National University, Seoul, South Korea

^40^Newcastle University, Newcastle, UK

^41^RWTH Aachen University, Aachen, Germany

^42^Caltech, Pasadena, USA

^43^Rockefeller University, New York, USA

^44^Mount Holyoke College, South Hadley, USA

^45^National Hualien Senior High School, Hualien, Taiwan

^46^The Francis Crick Institute, London, UK

^47^Aware LLC, Zurich, Switzerland

^48^University of California, Los Angeles, USA

^49^Institute of Biology, Leipzig University, Leipzig, Germany

^50^University of California, Merced, USA

^51^University of Florida, Gainesville, USA

^52^Retired MD-PhD, Fort Collins, USA

## Acknowledgements

We thank John Wiggins, G. McGrath, and Dave Barlieb for computer system administration and M. Husseini for project administration. We are grateful to J. Maitin-Shepard for Neuroglancer. We thank Pedro Nunez Gomez for help with GPU-cluster deployment. We thank the management at SixEleven and Ariadne for coordination and proofreader management. Mala Murthy and Sebastian Seung acknowledge support from the National Institutes of Health (NIH) BRAIN Initiative RF1 MH117815, RF1 MH129268 and U24 NS126935, from the Princeton Neuroscience Institute, as well as assistance from Google. Davi Bock was supported by NIH NIMH BRAIN Initiative grant 1RF1MH120679-01 and a Neuronex2 award (NSF 2014862). Gregory S.X.E. Jefferis and Davi Bock were supported by Wellcome Trust Collaborative Award (203261/Z/16/Z). Gregory S.X.E. Jefferis was supported by Wellcome Trust Collaborative Award 220343/Z/20/Z, Neuronex2 award (MRC MC_EX_MR/T046279/1) and received core support from the MRC (MC-U105188491). Albert Lin was supported by the NSF through the Center for the Physics of Biological Function (PHY-1734030). Ibrahim Tastekin was supported with a Marie Skłodowska-Curie postdoctoral fellowship (H2020-WF-01-2018-867459 to Ibrahim Tastekin) and by the Portuguese Research Council (Grant PTDC/MED-NEU/4001/2021). Andrew Seeds and Stefanie Hampel were supported by National Institute Of Neurological Disorders And Stroke of the National Institutes of Health under Award Number RF1NS121911. Derrick Brittain, Casey Schneider-Mizell, and Forrest Collman thank the Allen Institute for Brain Science founder, P. G. Allen, for his vision, encouragement and support. This work was also supported by the Intelligence Advanced Research Projects Activity via Department of Interior/Interior Business Center contract no. D16PC0005 to H.S.S. The US Government is authorized to reproduce and distribute reprints for Governmental purposes notwithstanding any copyright annotation thereon. The views and conclusions contained herein are those of the authors and should not be interpreted as necessarily representing the official policies or endorsements, either expressed or implied, of Intelligence Advanced Research Projects Activity, Department of Interior/Interior Business Center or the US Government.

## Contributions

Members of the FlyWire consortium contributed proofreading and annotations (see Supplemental Tables 1, 2). SGerhard provided braincircuits.io. TM and NK realigned the dataset with methods developed by EM, BN and TM and infrastructure developed by SP, ZJ. JAB, SM wrote code for masking defects and misalignments. KL trained the convolutional net for boundary detection, using ground-truth data realigned by DI. JW used the convolutional net to generate an affinity map that was segmented by RL. NK, MAC, OO, AH, CSJ, KKuehner and ARS adapted and improved Neuroglancer for proofreading and annotations. JG, KKruk, AM, SD, FC and CSM created interactive analysis and annotation tools for the community. AM created Codex with help from ARS, SD, KKuehner and RM. ARS and AM created the website. ARS, CEM and MS onboarded community members and tested new proofreaders. ARS, MS, CSJ and CEM designed tutorials. CEM, ARS and MS provided community support. SD, FC, CSM, CSJ, AH, DBrittain and WMS built and maintained CAVE for FlyWire and managed user access. SD, PS, AM and EP curated the data and made it available for download. EP and DDB provided a coordinate mapping service. ASB, NE, GSXEJ and JF provided neurotransmitter information. SCY, CEM, MC, KE, YY and PS trained and managed proofreaders. SD, SCY, PS and GSXEJ led the targeted proofreading effort. SD, PS, AM, AChampion and KKuehner maintained the proofreading management platforms. SD evaluated the proofreading accuracy. SD, AL, HSS, DD and RY analyzed the data. SD, DBland and SCY annotated and analyzed the ocellar circuit. SD, HSS, MM, AL, PS and ARS wrote the manuscript with feedback from ASB, WHuetteroth, GSXEJ and contributions from all authors. HSS, MM, GSXEJ, DDB sponsored large-scale proofreading. GSXEJ, DDB led the Cambridge effort. MM, HSS led the overall effort.

## Methods

### Neuropils

Meshes for individual neuropils were based on work by Ito et al.^110^. More specifically, we took meshes previously generated from a full brain segmentation of the JFRC2 template brain which are also used by the Virtual Fly Brain project (see also https://natverse.org/nat.flybrains/reference/JFRC2NP.surf.html). These meshes were moved from JFRC2 into FlyWire (FAFB14.1) space through a series of non-rigid transforms. In addition, we also generated two neuropil meshes for the laminae and for the ocellar ganglion. For these, the FlyWire synapse cloud was voxelized with 2 µm isotropic resolution, meshed using the marching cube algorithm using Python and manually post-processed in Blender 3d.

We calculated a volume for each neuropil using its mesh. In the aggregated volumes presented in the paper we assigned the lamina, medulla, accessory medulla, lobula, lobula plate to the optic lobe. The remaining neuropils but the ocellar ganglion were assigned to the central brain.

### Neuropil synapse assignments

We assigned synapses to neuropils based on their presynaptic location. We used ncollpyde (https://pypi.org/project/ncollpyde/) to calculate if the location was within a neuropil mesh and assigned the synapse accordingly. Some synapses remained unassigned after this step because the neuropils only resemble rough outlines of the underlying data. We then assigned all remaining synapses to the closest neuropil if the synapse was within 10 µm from it. The remaining synapses were left unassigned.

### Correction of left-right inversion

Our reconstruction used the FAFB EM dataset^52^. A number of consortium members (A. Bates, P. Kandimalla, S. Noselli) alerted us that the FAFB imagery seemed left-right inverted based on cell types innervating the asymmetric body^164^. Eventually a left-right inversion during FAFB imaging was confirmed. All side annotations in figures, in Codex and elsewhere are based on the true biological side. For technical reasons we were unable to invert the underlying FAFB image data and therefore continue to show images and reconstructions in the same orientation as^52^ although we now know in such frontal views the fly’s left is on the viewer’s left. For full details of this issue including approaches to display FAFB and other brain data with the correct chirality, please see our companion paper^44^.

### Proofreading system

FlyWire uses the Connectome Annotation Versioning Engine (CAVE) for hosting the proofreadable segmentation and all of its annotations. CAVE’s proofreading system is the PyChunkedGraph which has been described in detail elsewhere^55,165^.

### Proofreading annotations

Any user in FlyWire was able to mark a cell as complete, indicating that a cell was good for analysis. However, such annotations did not prevent future proofreading of a cell as commonly smaller branches were added later on. We created an annotation table for these completion markings. Each completion marking was defined by a point in space and the cell segment that overlapped with this point at any given time during proofreading was associated with the annotation. We created a webservice allowing users to submit completion markings for any cell. For convenience, we added an interface to this surface directly into Neuroglancer such that users can submit completion information for cells right after proofreading (Supplemental Information 1). When users submitted completion annotations we also recorded the current state of the cell. We encouraged users to submit new completion markings for a cell that they edited to indicate that edits were intentional. Recording the status of a cell at submission allowed us to calculate volumetric changes to a cell through further proofreading and flag cells for review if they received substantial changes without new completion markings.

### Onboarding proofreaders

Proofreaders came from several distinct labor pools: community members, citizen scientists from Eyewire (Flyers), and professional proofreading teams at Princeton and Cambridge. Proofreaders at Princeton consisted of staff at Princeton University and at SixEleven. Similarly, proofreading at Cambridge was performed by staff at Cambridge University and Ariadne. All proofreaders completed the built-in interactive tutorial and directed to Self-Guided Proofreading Training. For practice and learning purposes, the Sandbox, a complete replica of the FlyWire data, allowed new users to freely make edits and explore without affecting the actual “Production” dataset. When ready, an Onboarding Coordinator tested the new proofreader before giving access to the Production dataset^55^. Later onboarding called for users to send demonstration Sandbox edits that were reviewed by the Onboarding Coordinator. A new class of view-only users was introduced in early 2023, allowing researchers early data access for analysis purposes. All early access users attended a live onboarding session in Zoom prior to being granted edit or view access.

### Training the professional proofreading team

The professional proofreading team received additional proofreading training. Correct proofreading relies on a diverse array of 2D and 3D visual cues. Proofreaders learned about 3D morphology, resulting from false merger or false split without the knowledge of knowing what types of cells they are. Proofreaders studied various types of ultrastructures as the ultrastructures provide valuable 2D cues and serve as reliable guides for accurate tracing. Before professional proofreaders were admitted into Production, each of them practiced on average >200 cells in a testing dataset where additional feedback was given. In this dataset, we determined the accuracy of test cells by comparing them to ground-truth reconstructions. To improve proofreading quality, peer learning was highly encouraged.

### Recruitment of citizen scientists

The top 100 players from Eyewire, a gamified EM reconstruction platform that crowdsources reconstructions in mouse retina and zebrafish hindbrain^58^, received an invitation to beta test proofreading in FlyWire. A new set of user onboarding and training materials were created for citizen scientists, including: a blog, forum, and public Google docs. We created bite-sized introduction videos, a comprehensive “FlyWire 101” resource, as well as an Optic Lobe Cell Guide to aid users in understanding the unique morphology of flies. A virtual Citizen Science Symposium introduced players to the project, after which the self-dubbed “Flyers” began creating their own resources, such as a new comprehensive visual guide to cell types, conducting literature reviews, and even developing helpful FlyWire plugins. As of publication, FlyWire has 12 add-on apps ranging from a batch processor to cell naming helper (https://blog.flywire.ai/2022/08/11/flywire-addons/).

### Proofreading strategy to complete the connectome

As previously described^55^, proofreading of the connectome was focused on the microtubule-rich ‘backbones’ of neurons. Microtubule-free ‘twigs’ were only added if discovered incidentally or sought out specifically by members of the community. After proofreading, users marked neuronal segments as ‘complete’ indicating that neurons were ready for analysis but further changes remained possible. While *Drosophila* neuroscientist members of the FlyWire community generally contributed proofreading for their neurons of interest, the bulk of the segments was proofread by professional proofreaders in the following way: first we proofread all segments with an automatically detected nucleus in the central brain^81^ by extending it as much as possible and removing all false mergers (pieces of other neurons or glia attached), and second, going through the remaining segments in descending order of their synapse count (pre+post) up to a predefined size threshold of 100 synapses.

### Quality Assurance

To assess quality, a group of expert centralized proofreaders conducted a review of 3106 segments in the central brain. These specific neurons were chosen based on certain criteria such as significant change since being marked complete and small overall volume. An additional 826 random neurons were included in the review pool as well. Proofreaders were unaware which neurons were added for quality measurement and which ones because they were flagged by a metric. We compared the 826 neurons before and after the review and found that the initial reconstruction scored an average F1-Score of 99.2% by volume (Ext. Data Fig. 1-2a,b).

### Quantification of proofreading effort

Any quantification of the total proofreading time that was required to create the FlyWire resource is a rough estimate because of the distributed nature of the community, the interlacing of analysis and proofreading and the variability in how proofreading was performed. The first public release, version 630, required 2,712,769 edits. We measured proofreading times during early proofreading rounds that included proofreading of whole cells in the central brain. We collected timings and number of edits for 29,135 independent proofreading tasks after removing outliers with more than 500 edits. From this data we were able to calculate an average time per edit. However, we observed that proofreading times per edit were much higher for proofreading tasks that required few edits (<5). That meant that our measurements were not representative for the second round of proofreading which went over segments with > 100 synapses. These usually required 1-5 edits. We adjusted for that by computing estimates for proofreading speeds of both rounds by limiting the calculations to a subset of the timed tasks: (round 1) The average time per edit in our proofreading time dataset, (round 2) the average time of tasks with 1-5 edits. We average these times for an overall proofreading time because the number of tasks in each category were similar. The result was an average time of 79s per edit which adds up to an estimate of 29.8 person-years assuming a 2000h work year.

### Completion rates

We adopted the completion rate calculations from the hemibrain^33^. Every presynaptic and postsynaptic location was assigned to a segment. Using the neuropil assignments, we then calculated the fraction of presynapses that were assigned to segments marked as proofread for each neuropil and analogous for postsynaptic location.

### Comparison with the hemibrain

We retrieved the latest completion rates and synapse numbers for the hemibrain from neuprint (v1.2.1). In some cases, neuropil comparisons were not directly possible because of redefined regions in the hemibrain dataset. We excluded these regions from the comparison.

### Crowdsourced annotation

FlyWire’s large community and diversity of expertise allowed us to crowdsource the identification of neurons. There is no limit to the number of annotations a neuron can receive. A standardized format is encouraged but not required. One user might first report that a neuron is a descending interneuron, while another might add that it is the Giant Fiber descending neuron, and another might add all its synonyms and citations from the literature. Contributors’ names are visible so they can be consulted if there is disagreement. The disadvantage to this approach is that there isn’t one precise name for every neuron, but the advantage is a richness of information and dialog. The annotations are not meant to be a finished, static list, but a continually growing, living data source. These annotations were solicited from the FlyWire community through Town Halls, email announcements, interest groups in the FlyWire Forum, online instructions, and by personal contact from the Community Manager. Citizen scientists also contributed annotations, after receiving training on particular cell types by experts.

### Neuron categorizations

Neuron categorization, sensory modality annotations and nerve assignments are described in detail in our companion paper^121^ In brief, neurons were assigned to one of three “flow” classes: afferent (to the brain), intrinsic (within the brain), and efferent (out of the brain). Intrinsic neurons had their entire arbor within the FlyWire dataset. This included cells that projected to and from the subesophageal zone (SEZ). Next, each flow class was divided into “super” classes in the following way. afferent: sensory, ascending. intrinsic: central, optic, visual projection (from the optic lobes to the central brain), visual centrifugal (from the central brain to the optic lobes). efferent: endocrine, descending, motor.

### Skeletonization and path length calculation

We generated skeletons for all neurons marked as proofread using skeletor (https://github.com/navis-org/skeletor) which implements multiple skeletonization algorithms such as TEASAR^166^. In brief, neuron meshes from the exported segmentation (LOD 1) were downloaded and skeletonized using the “wavefront” method in skeletor. These raw skeletons were then further processed (e.g. to remove false twigs and heal breaks) and produce downsampled versions using navis (https://github.com/navis-org/navis). A modified version of this skeletonization pipeline is implemented in fafbseg (https://github.com/navis-org/fafbseg-py).

### Synaptic connections

We imported the automatically predicted synapses from Buhmann et al.^45^ which we combined with the predictions by Heinrich et al. to assign scores to all synapses ^46^ to improve precision. We removed synapses from the imported list if they fulfilled any of the following criteria: (1) either the pre- or postsynaptic location remained unassigned to a segment (proofread or unproofread), (2) It had a score ≤50.

### Connection threshold

For all the analyses presented in this paper, save for synapse distributions, we employed a consistent threshold of >4. Our decision to use a synapse threshold on connections was due partly to the fact that synapses in the FlyWire dataset were not manually proofread. For these analyses, many of which demonstrate the high interconnectivity of the fly brain, we chose a conservative threshold to ensure that considered connections are real. Use of a threshold is also in keeping with previous work analyzing wiring diagrams in *Drosophila* ^33^. Thus, we are likely undercounting the number of true connections. The distribution of synapse counts (Fig. 3f) does not display any bimodality that could be used to set the threshold. Therefore, the choice of 5 synapses per connection is a reasonable but arbitrary one. In the companion paper analyzing the network properties of the FlyWire connectome, it is found that statistical properties of the whole-brain network, such as reciprocity and clustering coefficient, are robust to our choice of threshold (Lin et al., *in prep*). The FlyWire data is available without an imposed threshold, so users can choose their own appropriate threshold for their specific use case.

### Neuropil projectome construction

Under the simplifying assumptions that information flow through the neuron can be approximated by the fraction of synapses in a given region, and that inputs and outputs can be treated independently, we can construct a matrix representing the projections of a single neuron between neuropils. The fractional inputs of a given neuron are a 1 x N vector containing the fraction of incoming synapses the neuron has in each of the N neuropils, and the fractional outputs are a similar vector containing the fraction of outgoing synapses in each of the N neuropils. We multiply these vectors against each other to generate the N x N matrix of the neuron’s fractional weights. Summing these matrices across all intrinsic neurons produces a matrix of neuropil-to-neuropil connectivity (Fig. 4a). In this projectome, all neurons contribute an equal total weight of one.

### Dominant input side

We assigned neuropils to the left and right hemispheres or the center if the neuropil has no homologue. We then counted how many postsynapses each neuron had in each of these three regions and assigned it to the one with the largest count.

### Contralateral and bilateral neuron analysis

For each neuron, we calculated the fraction of presynapses in the left and right hemisphere. The hemisphere opposite its dominant input side was named the contralateral hemisphere. We excluded neurons that had either most of their presynapses or most of their postsynapses in the center region.

### Rank analysis & Information Flow

We used the information flow algorithm implemented by Schlegel et al.^20^ (https://github.com/navis-org/navis) to calculate a rank for each neuron starting with a set of seed neurons. The algorithm traverses the synapse graph of neurons probabilistically. The likelihood of a neuron being added to the traversed set increased linearly with the fraction of synapses it receives from already traversed neurons up to 30% and was guaranteed above this threshold. We repeated the rank calculation for all sets of afferent neurons as seed as well as the whole set of sensory neurons. The groups we used are:

olfactory receptor neurons, gustatory receptor neurons, mechanosensory Johnston’s Organ neurons, head and neck bristle mechanosensory neurons, mechanosensory taste peg neurons, thermosensory neurons, hygrosensory neurons, visual projection neurons, visual photoreceptors, ocellar photoreceptors and ascending neurons.

Additionally, we created input seeds by combining all listed modalities, all sensory modalities, and all listed modalities with visual sensory groups excluded.

For each modality we then ordered the neurons according to their rank and assigned them a percentile based on their location in the order. To compute a reduced dimensionality, we treated the vector of all ranks (one for each modality) as neuron embedding and calculated two dimensional embeddings using UMAP^167^ with the following parameters: n_components=2, min_dist=0.35, metric=“cosine”, n_neighbors=50, learning_rate=.1, n_epochs=1000.

## Extended Data Figures

**Ext.-Figure 1-1.**
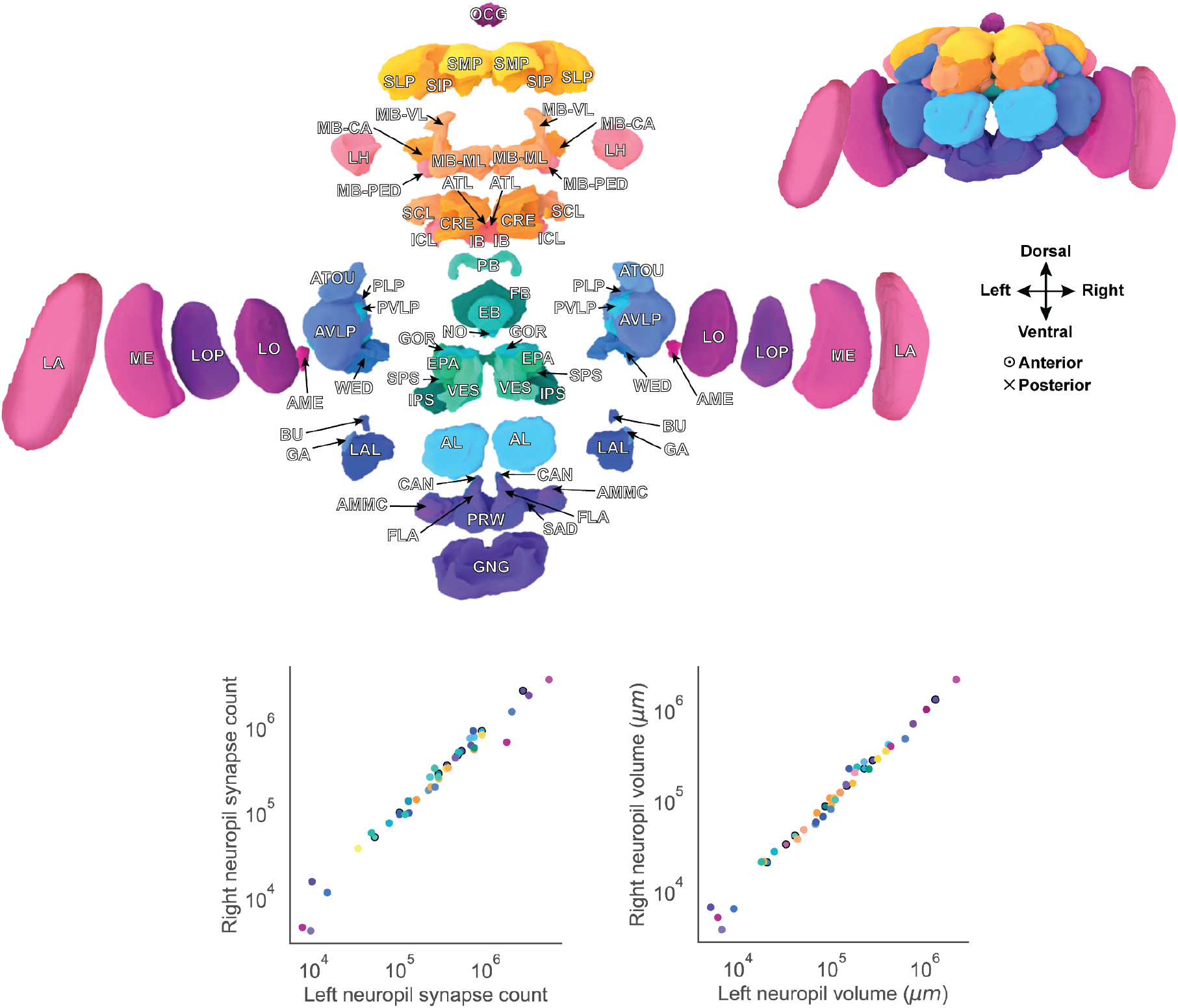
Neuropils of the fly brain.

**Ext. Figure 1-2.**
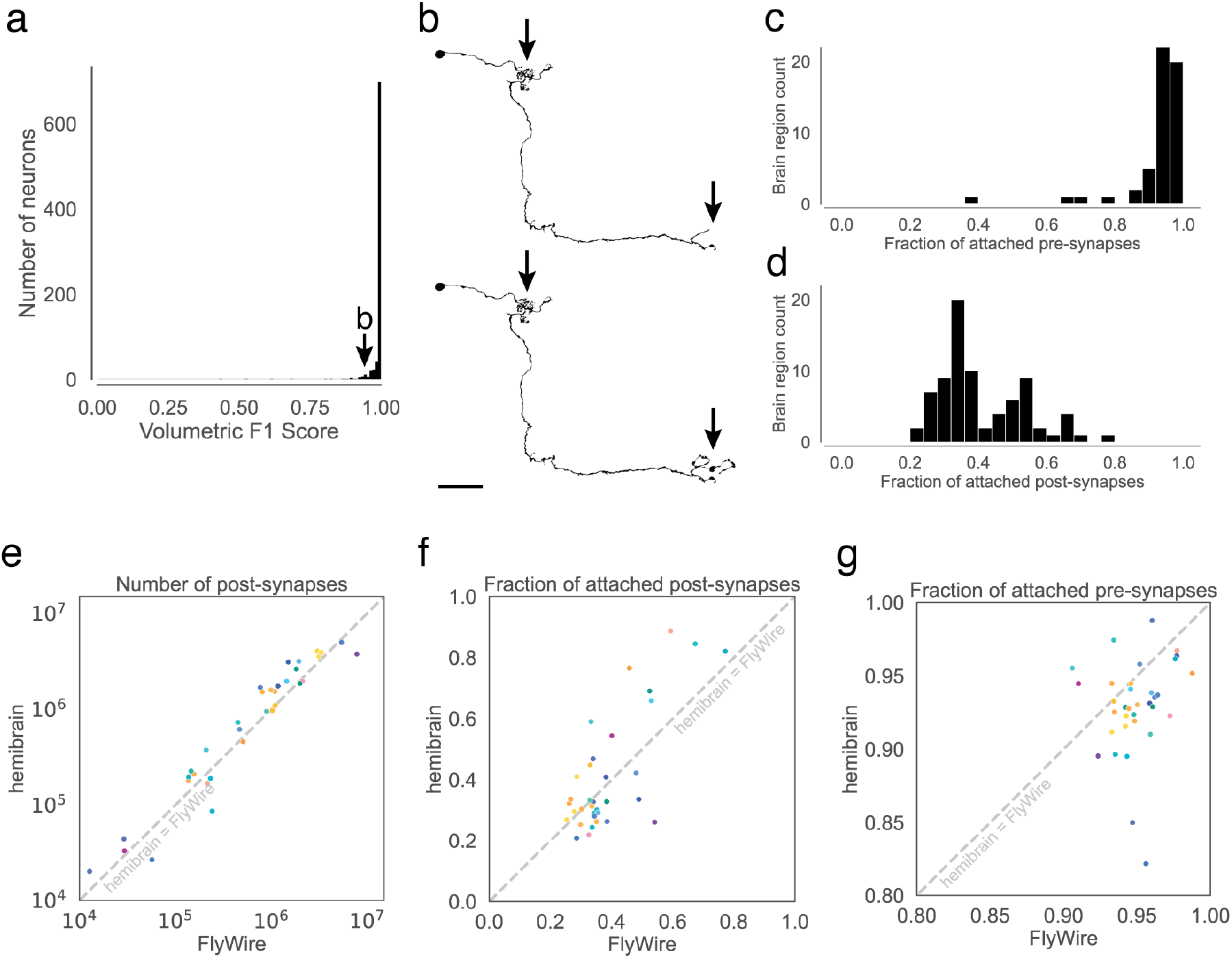
Completeness and accuracy of FlyWire’s reconstruction. (a) shows the result of our evaluation of proofread segments in the central brain. Experts attempted further proofreading of 826 neurons. We computed volumetric overlaps between the original and the final segment to calculate precision, recall, and F1 Scores. (b) Examples (top: before, bottom: after) of the changes made during further proofreading for a neuron scoring an F1-Score of 0.936. Arrows highlight locations that changed. (c,d) For each neuropil, we quantified what fraction of the synapses within it are pre- and postsynaptically attached to a proofread segment. (c) displays the distribution for presynaptic attachment and (d) the distribution for postsynaptic attachment. (e, f, g) Comparisons between FlyWire’s reconstruction and the hemibrain were made for overlapping neuropils. Dots represent neuropils and are colored according to Ext. Data Fig. 1-1. (e) Comparison of the number of automatically detected synapses. The axes are log-transformed. (f) Comparison of post-synaptic completion rates and (g) pre-synaptic completion rate. The axes are truncated.

**Ext. Figure 1-3.**
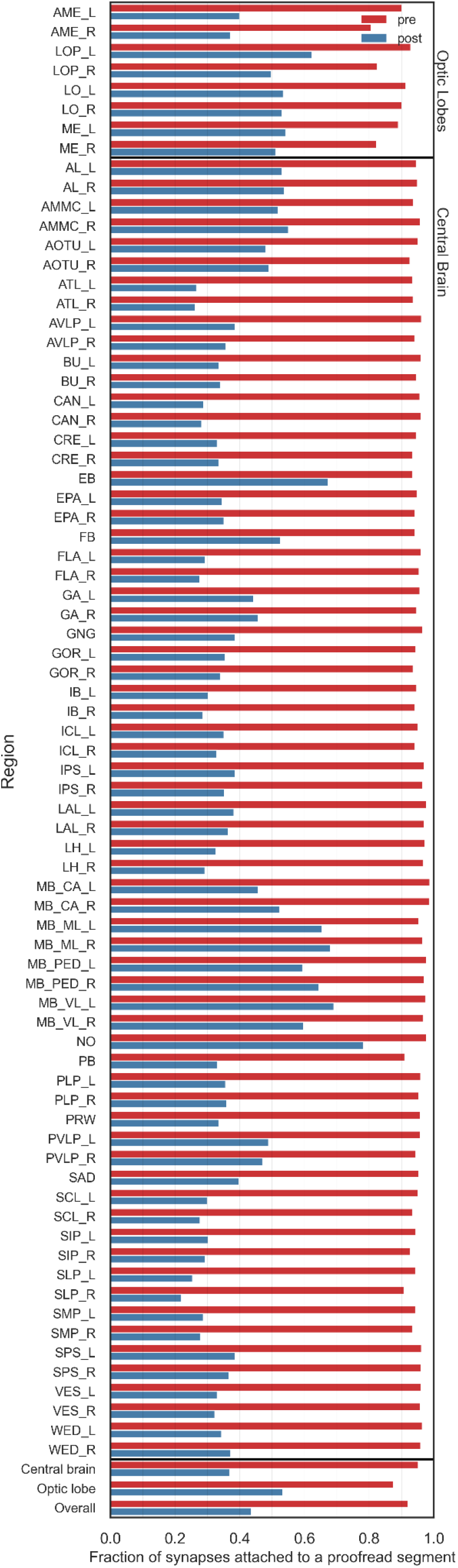
Completion rates by neuropil.

**Ext. Figure 1-4.**
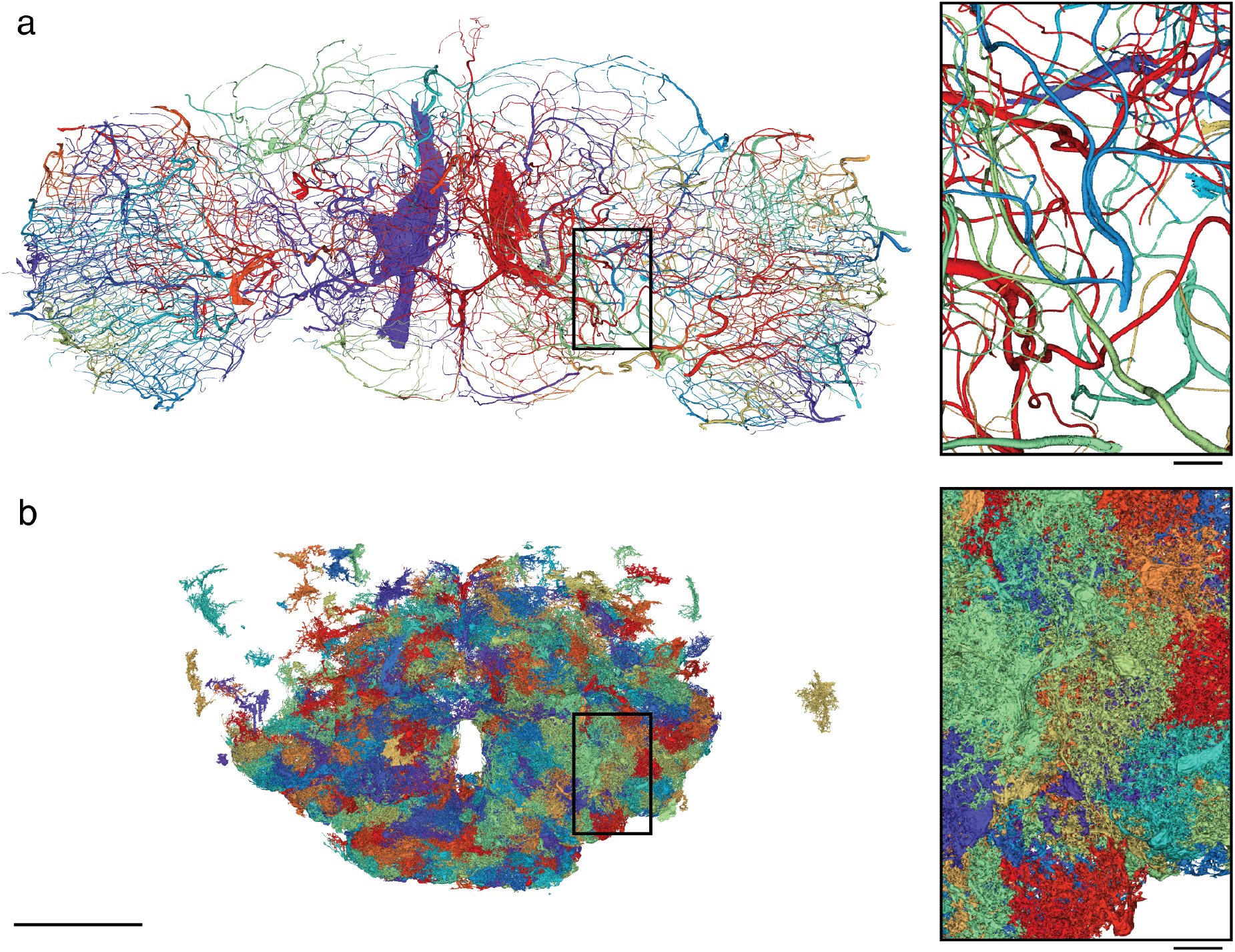
Trachea and glia cells. (a) Rendering of all trachea segments in the FlyWire dataset. (b) Rendering of some reconstructed glia cells in the FlyWire dataset. At the time of writing, only a subset of the glia cells, with bias towards the central brain, have been proofread and labeled. Scale bar: 100 µm; insets: 10 µm.

**Extended Data Figure 3-1.**
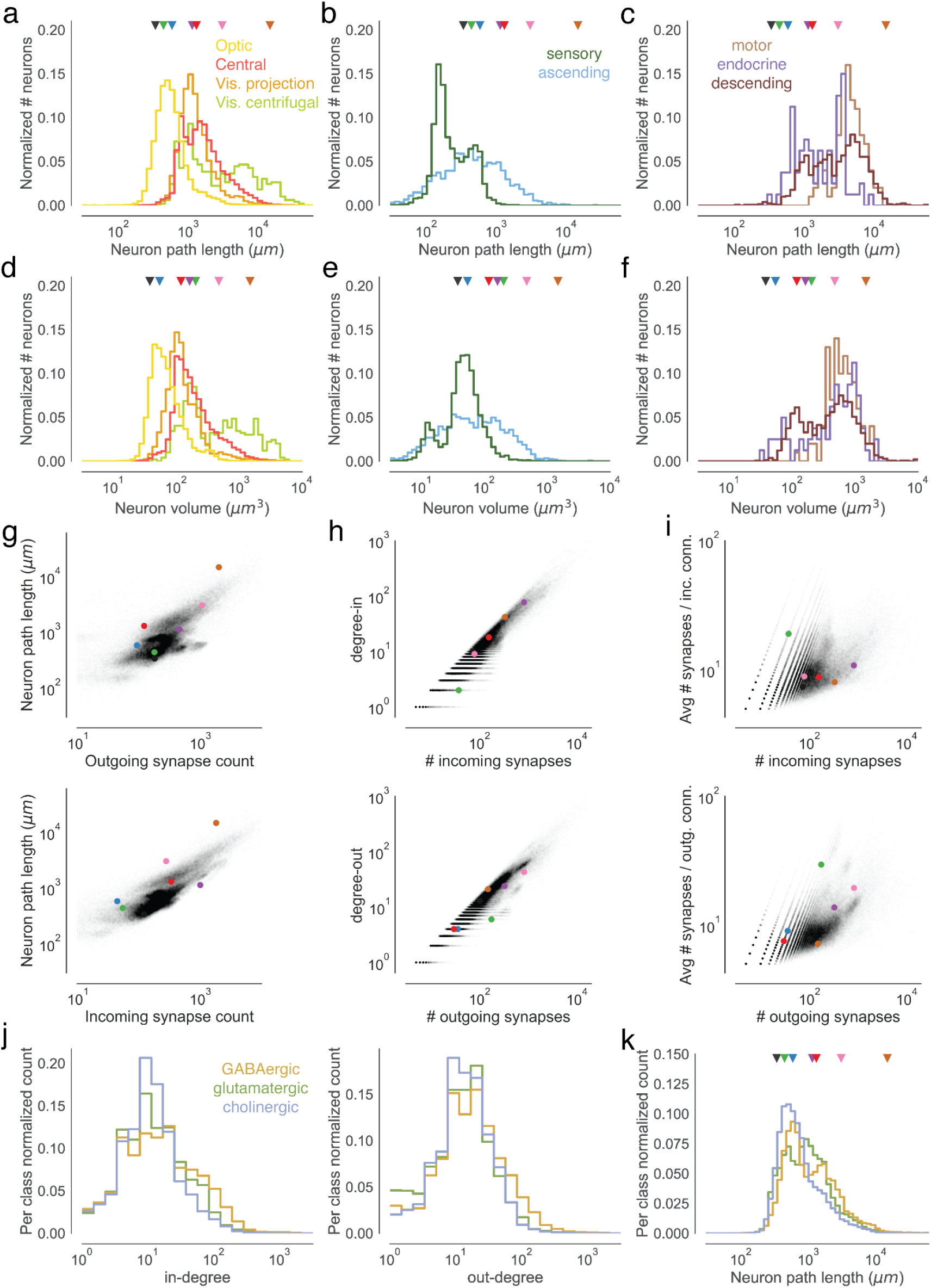
Measurements of neuron size. Colored markers refer to neurons in Fig. 3b. (a) Neuron path lengths of intrinsic neurons, (b) afferent neurons, and (b) efferent neurons by super-class. (d) Volumes of intrinsic neurons, (e) afferent neurons, and (f) efferent neurons by super-class. (g) Comparisons of path lengths and number of incoming and outgoing synapses. (h) For intrinsic neurons, comparisons of the in- and out-degrees with the number of incoming and outgoing synapses. Every dot is a neuron. (i) Comparison of average connection strengths (synapses per connection) with the number of synapses. Every dot is a neuron. (j) In- and out-degree distributions by neurotransmitter type. (k) Neuron path lengths by neurotransmitter type.

**Ext. Figure 4-1.**
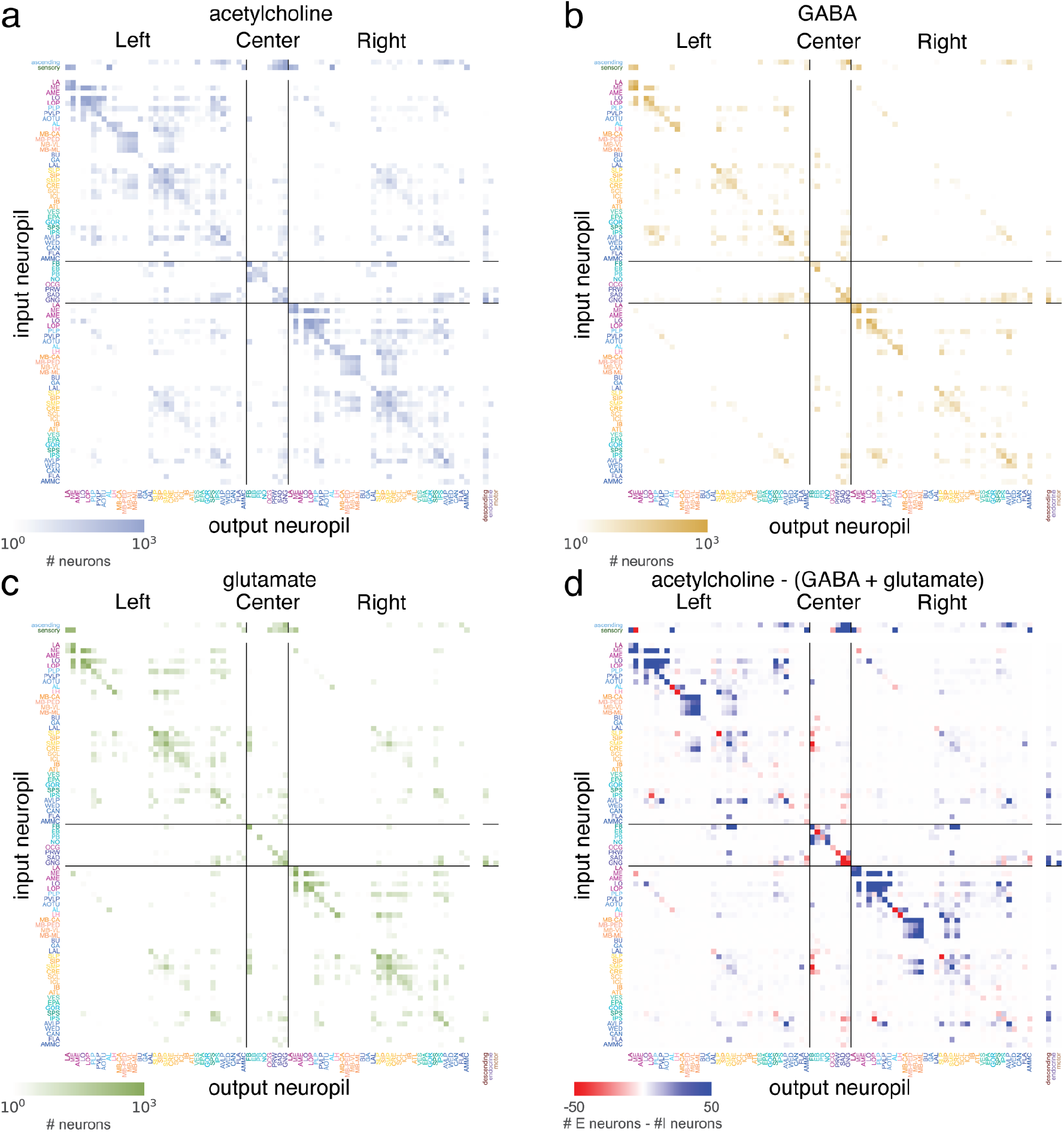
Neuropil-neuropil projection maps. (a) Projection maps produced as in Fig. 4a limited to connections from cholinergic, (b) GABAergic, and (c) glutamatergic neurons. (d) The difference between the putative excitatory (acetylcholine) and the putative inhibitory (GABA, glutamate) projection maps.

**Ext. Figure 4-2.**
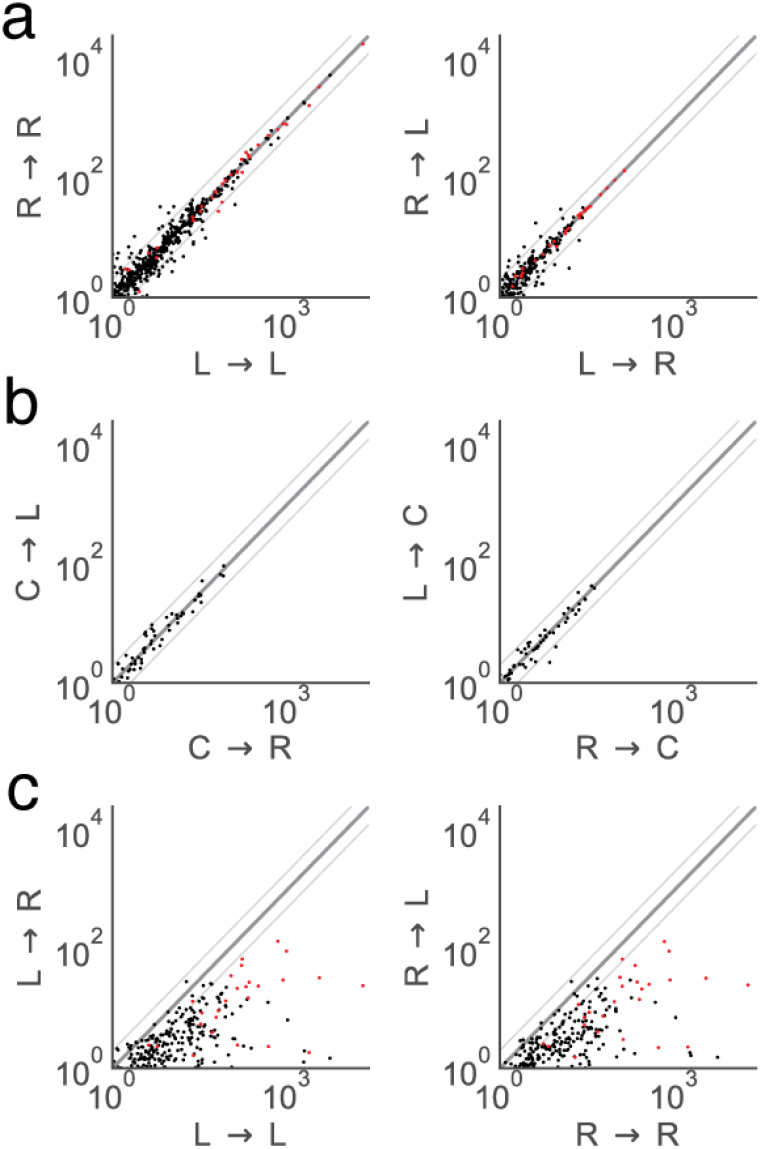
Neuropil-neuropil projections compared between hemispheres. Each dot is a neuropil-neuropil projection in one hemisphere and the axes show the fractional weights as calculated in Fig. 4a,b. Red dots are comparisons between the same neuropils in different hemispheres (e.g. AMMC(L) -> VLP(L) vs AMMC(R) -> VLP(R). (a) Comparison of projections between neuropils in both hemispheres and between hemispheres. (b) Comparisons of projections with the center neuropils. (c) Comparisons of projections between ipsilateral and contralateral neuropil projections.

**Ext. Figure 4-3.**
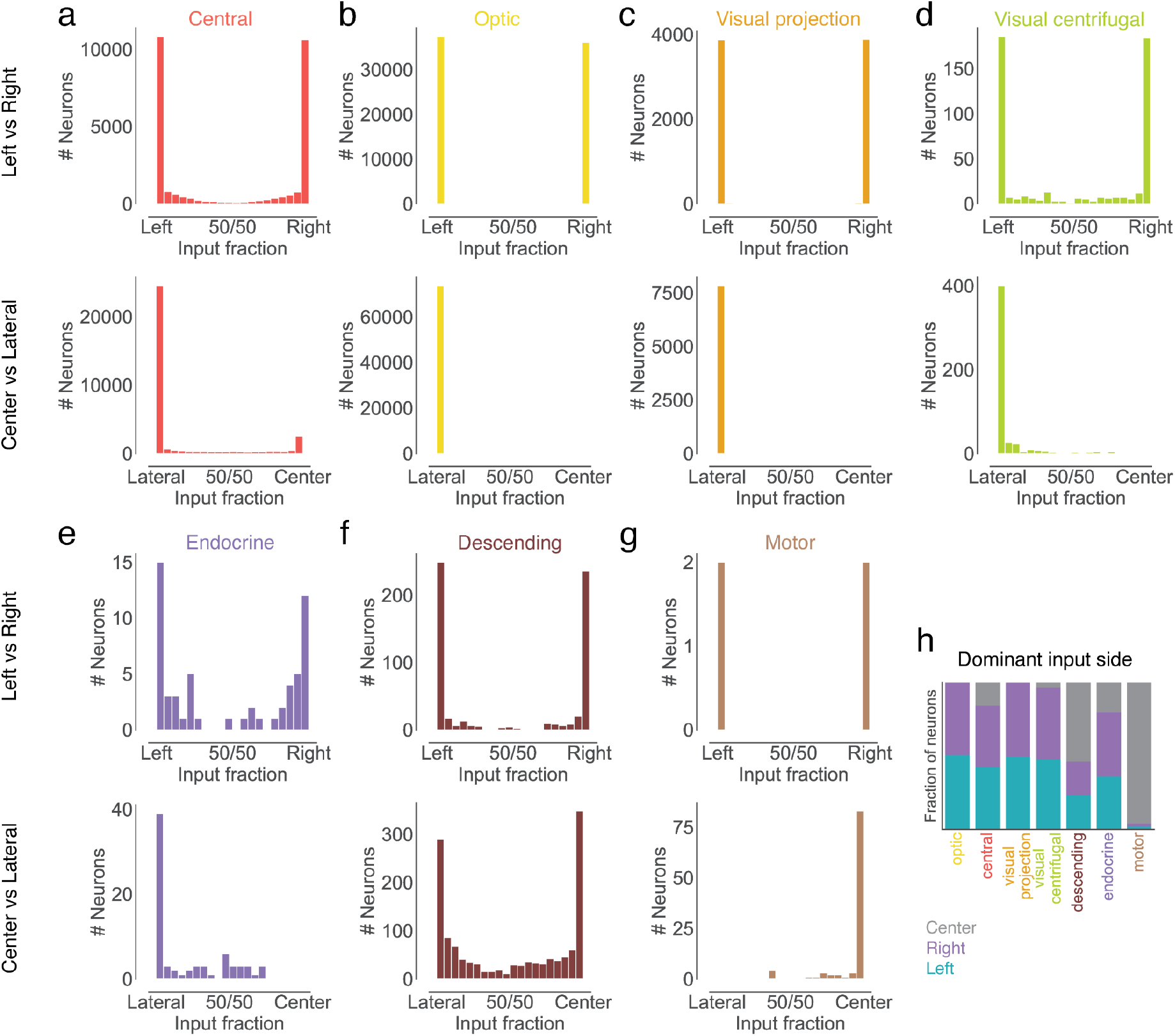
Input side analysis. We assigned postsynaptic locations to either the center region or the left or right hemisphere. (a-g) For each super-class, the fraction of synapses in the left vs right hemisphere is shown for those neurons receiving most of their neurons laterally (top plot). The lower plot shows the fraction of synapses in the center vs the lateral regions for all neurons. (h) Each neuron was assigned to the side where it received most of its inputs.

**Ext. Figure 4-4.**
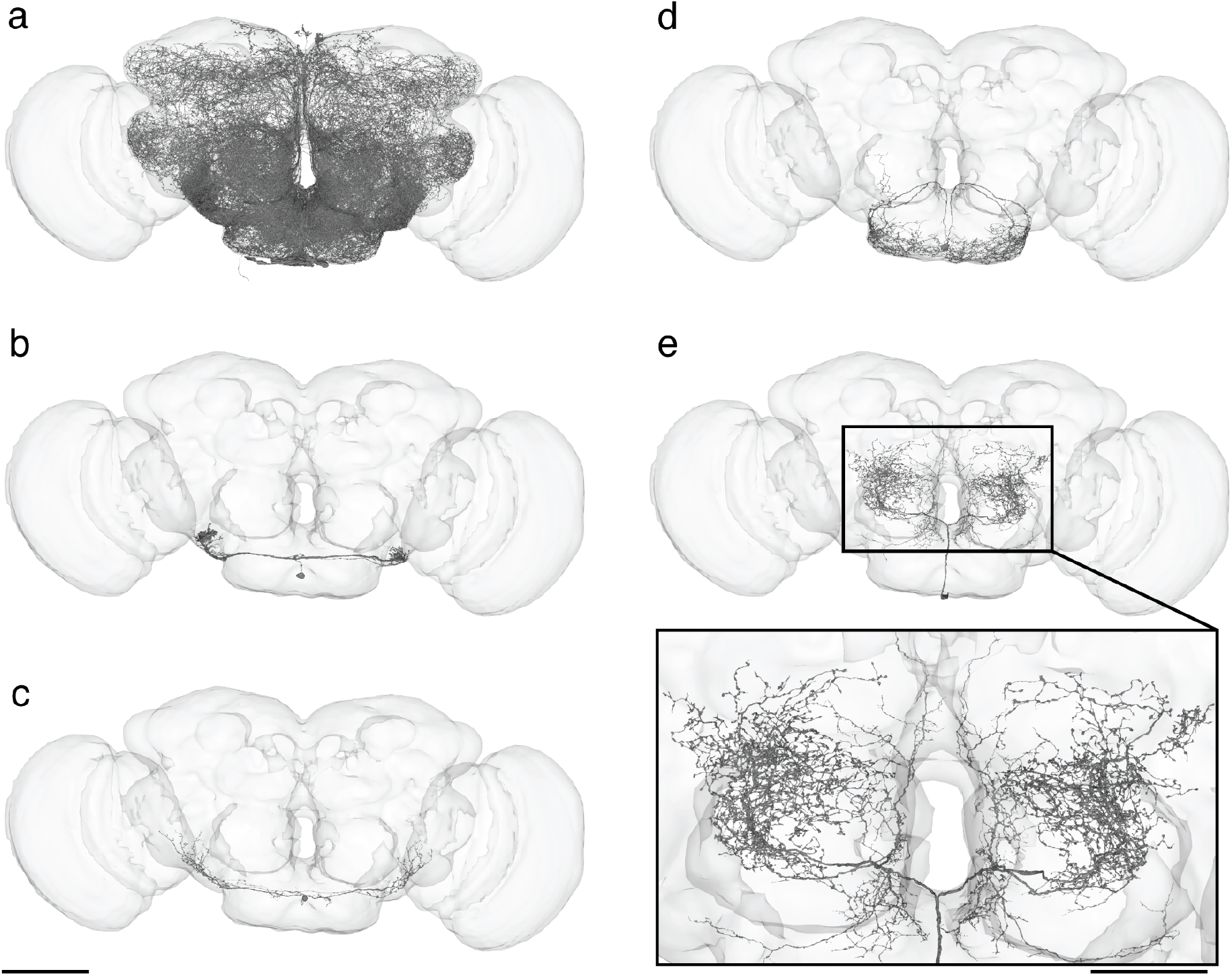
Neurons on the midline with dendrites in both hemispheres. (a) All symmetric neurons with a cell body on the midline (N=106). (b-e) examples of individual neurons. Scale bar: 100 µm, inset: 50 µm

**Ext. Figure 4-5.**
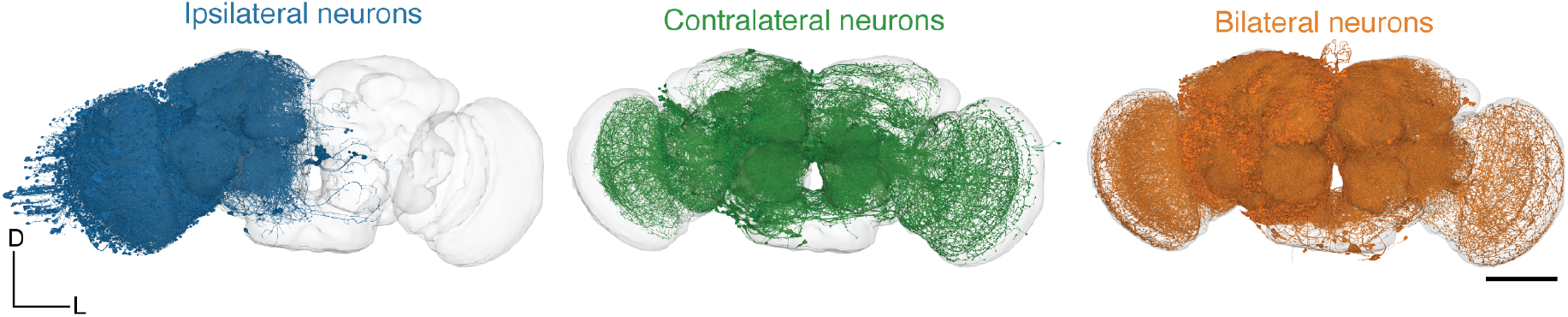
Renderings of neurons for each cross-hemisphere category. (up to 3,000 neurons rendered per group). Scale bar: 100 µm

**Extended Data Figure 6-1.**
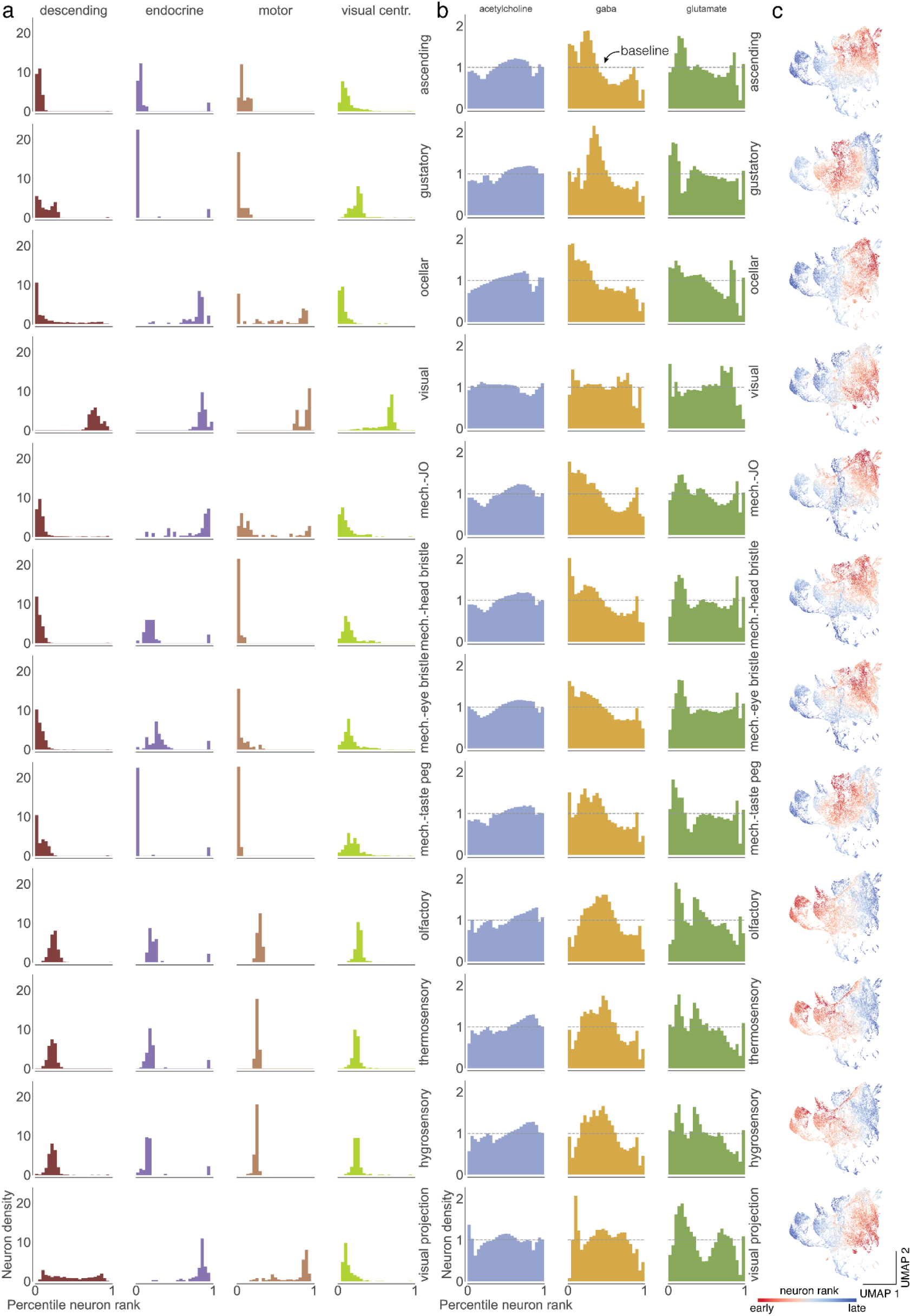
Percentile ranks for every modality. (a) For each sensory modality we used the traversal distances to establish a neuron ranking. Each panel shows the distributions of neurons of each super-class within the sensory modality specific rankings. (b) Same as in (a) for the fast neurotransmitters. (c) Neurons in the UMAP plot are colored by the rank order in which they are reached from a given seed neuron set. Red neurons are reached earlier than blue neurons.

**Extended Data Figure 6-2.**
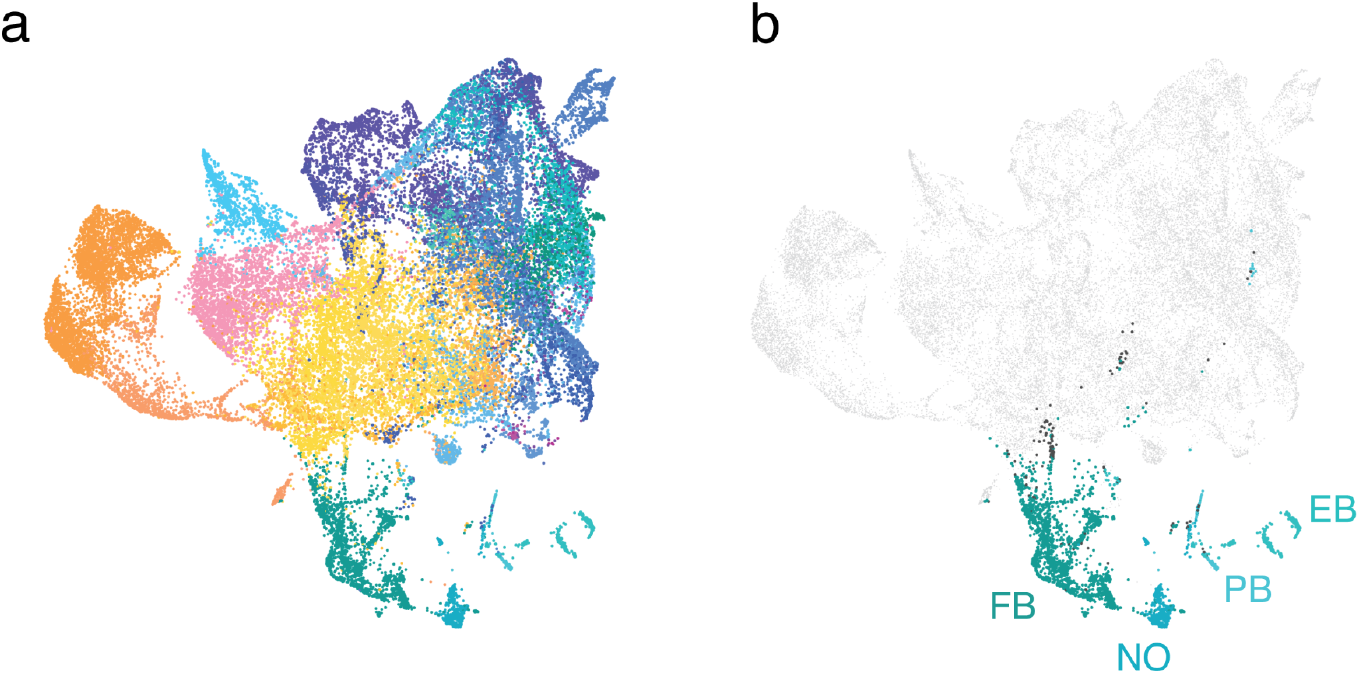
Rank-based UMAP projection and neuropils. (a) Every neuron in the central brain was assigned to the neuropil where it received the most synapses. Every dot is then colored by the assigned neuropil (Ext. Data Fig. 1-1). (b) Same as in a but limited to the central complex neurons. Neurons in the central complex with an assigned neuropil other than the ones shown are colored black.

**Extended Data Figure 7-1.**
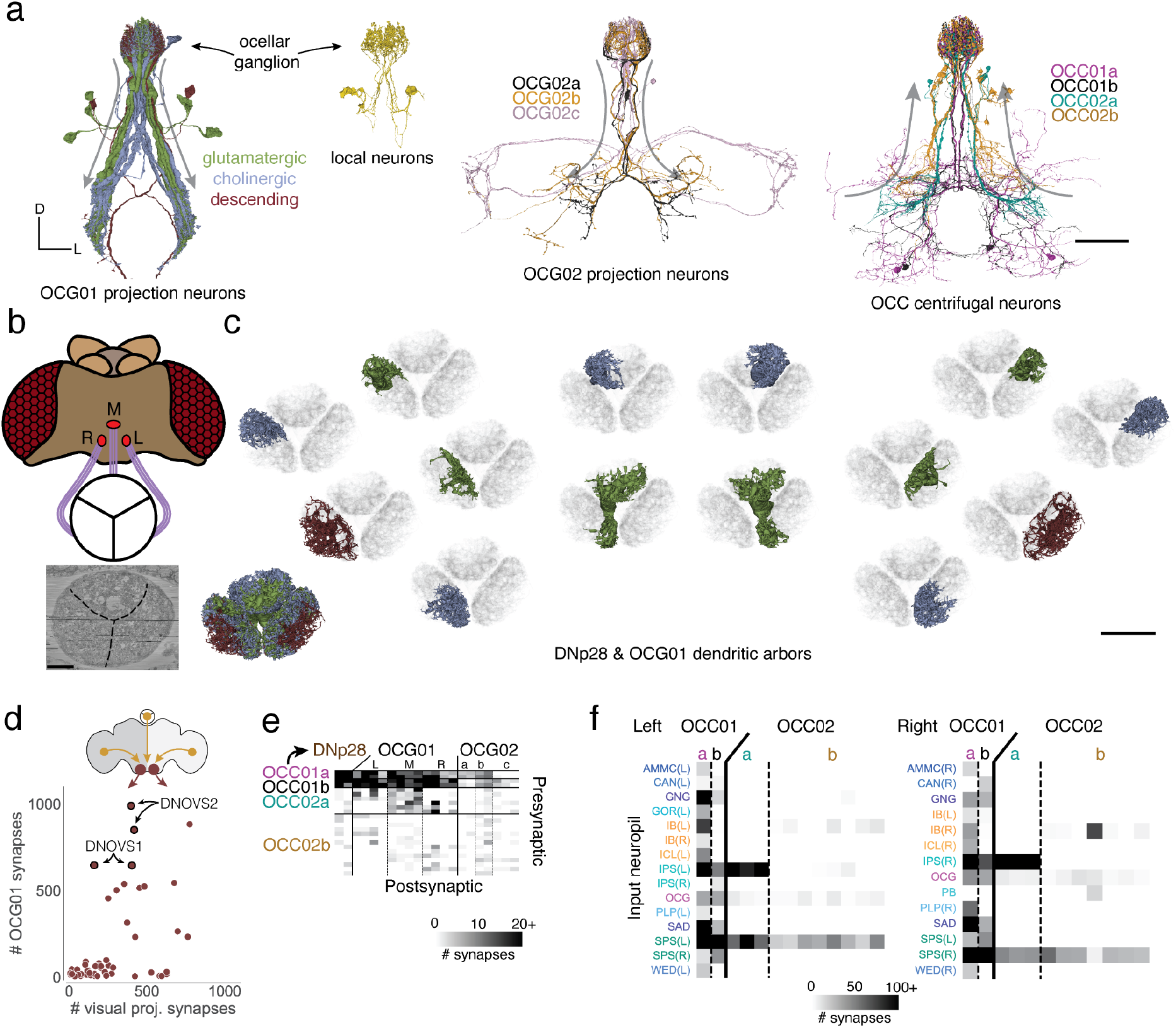
Ocellar circuit. (a) Renderings of all neurons (excluding the photoreceptors) with arbors in the ocellar ganglion. “Information flow” from pre- and postsynapses is indicated by arrows along the arbors. (b) Overview of the three ocelli (left, medial, right) which are positioned on the top of the head. Photoreceptors from each ocellus project to a specific subregion of the ocellar ganglion which are separated by glia (marked with black lines on the EM). (c) Top view of the dendritic arbors within the ocellar ganglion of each DNp28 (brown) and OCG01 (blue: cholinergic, green: glutamatergic). The render on the lower shows all 12 OCG01s and 2 DNp28s. Each other render shows one neuron in color and all others in the background in gray for reference. (d) Comparison of number of synapses from OCG01 neurons and visual projection neurons onto descending neurons. (e) Connectivity matrix for connections between ocellar centrifugal neurons and ocellar projection neurons. (f) Inputs to ocellar centrifugal neurons by neuropil. Scale bars: 100 µm (a), 20 µm (c)

## Supplementary Information

**Supplemental Information 1:**
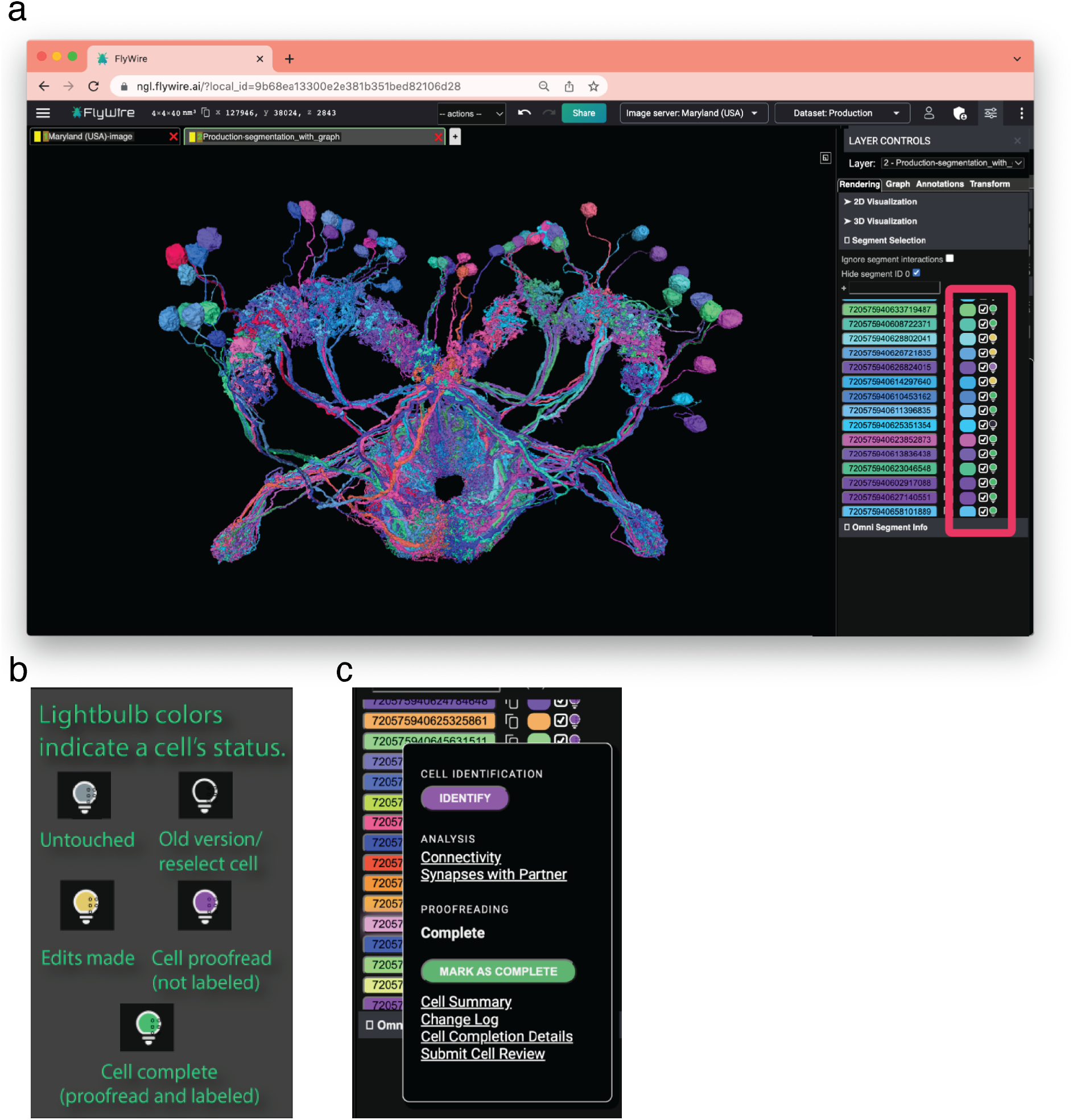
Neuroglancer Interface and Lightbulb. (a) FlyWire’s lightbulb menu displays the proofreading and annotation status of every segment (red box). (b) It is color coded for easy reference (yellow: cell has not been declared complete; purple: complete but not labeled; green: proofread and labeled; black: out of date segmentation). (c) Users can load cell identification directly within the FlyWire editor, perform basic connectivity analysis, and view a cell’s edit history.

**Supplemental Information 2:**
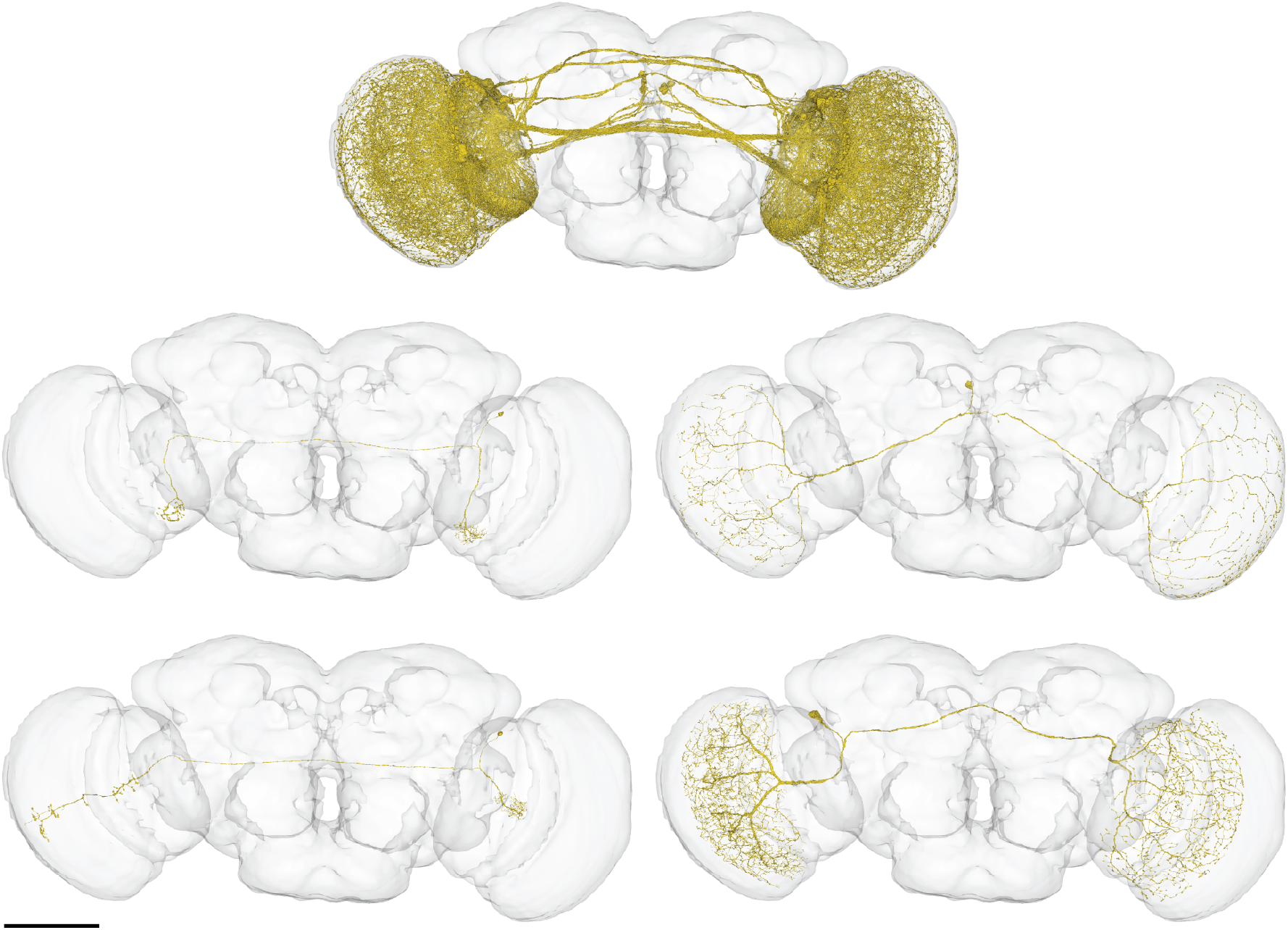
Bilateral optic lobe neurons. On the left: putative LC14 (top) and putative LC14b (bottom). Scale bar: 100 µm

**Supplemental Information 3:**
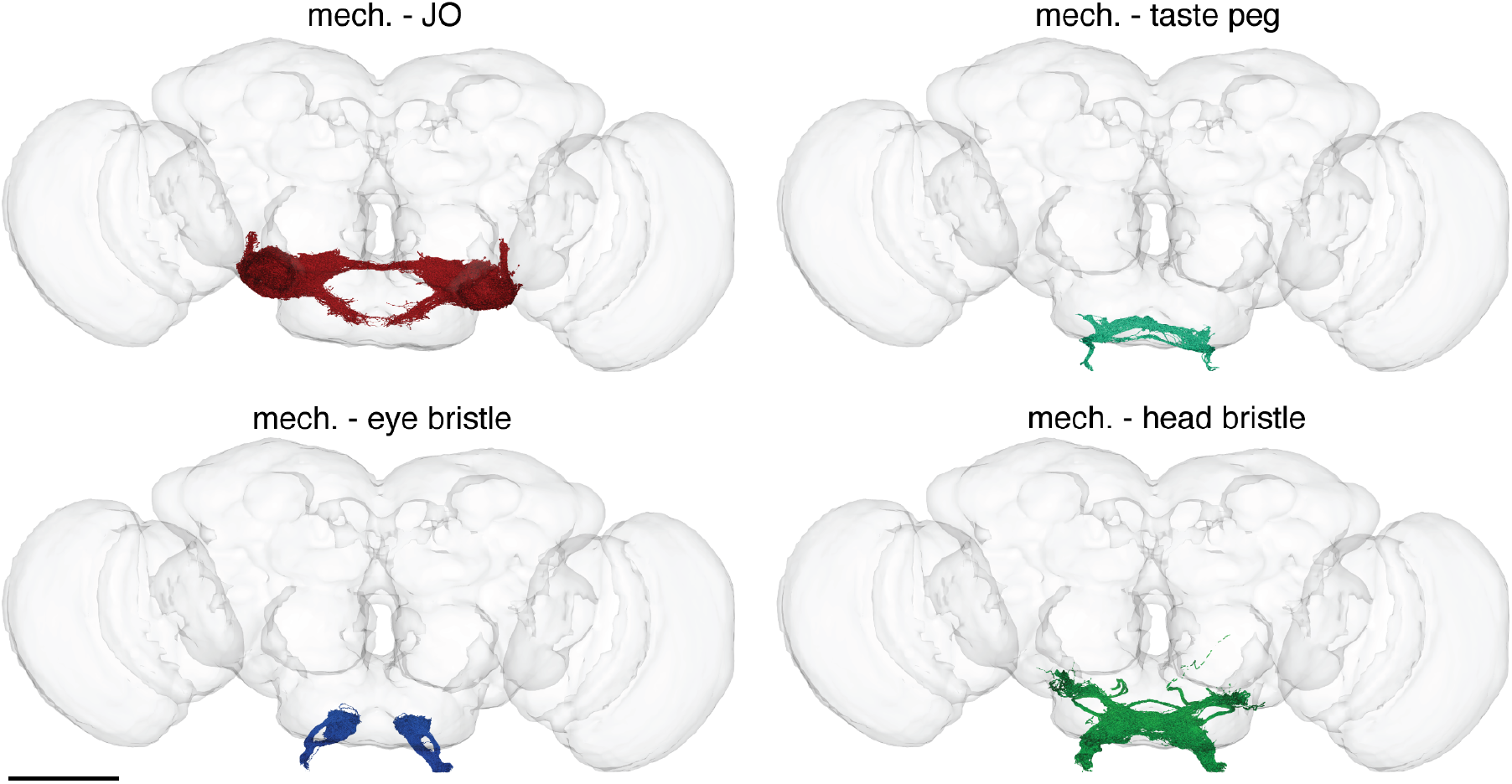
Mechanosensory neuron subtypes. Scale bar: 100 µm

**Supplemental Information 4:**
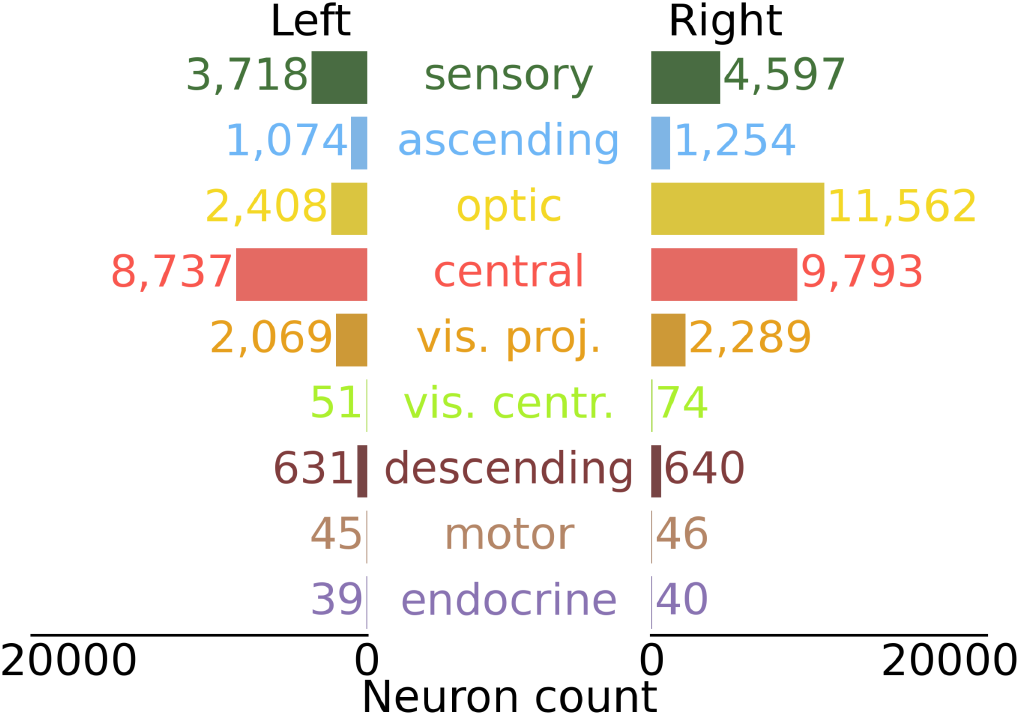
Distribution of community annotations by cell type.

**Supplementary Table 1:** Number of proofreading edits by consortium members. Only members with ≥100 edits are shown.

| Name | Lab Affiliation | total edits | specific contributions |
| --- | --- | --- | --- |
| Doug Bland | Mala Murthy Lab, Sebastian Seung Lab | 247,366 |  |
| Zairene Lenizo | Mala Murthy Lab, Sebastian Seung Lab | 122,932 |  |
| Nseraf | Flyers (citizen scientist) | 110,082 |  |
| Kyle Patrick Willie | Mala Murthy Lab, Sebastian Seung Lab | 83,635 |  |
| Nashra Hadjerol | Mala Murthy Lab, Sebastian Seung Lab | 81,700 |  |
| Austin T Burke | Mala Murthy Lab, Sebastian Seung Lab | 80,647 |  |
| Ryan Willie | Mala Murthy Lab, Sebastian Seung Lab | 74,951 |  |
| John Anthony Ocho | Mala Murthy Lab, Sebastian Seung Lab | 67,436 |  |
| Arti Yadav | Greg Jefferis Lab | 59,394 |  |
| Joshua Bañez | Mala Murthy Lab, Sebastian Seung Lab | 59,014 |  |
| Shirleyjoy Serona | Mala Murthy Lab, Sebastian Seung Lab | 55,859 |  |
| Yijie Yin | Greg Jefferis Lab | 50,805 |  |
| Rey Adrian Candilada | Mala Murthy Lab, Sebastian Seung Lab | 50,492 |  |
| Jet Ivan Dolorosa | Mala Murthy Lab, Sebastian Seung Lab | 46,454 |  |
| Mendell Lopez | Mala Murthy Lab, Sebastian Seung Lab | 43,885 |  |
| Ariel Dagohoy | Mala Murthy Lab, Sebastian Seung Lab | 43,124 |  |
| Regine Salem | Mala Murthy Lab, Sebastian Seung Lab | 42,181 |  |
| Griffin Badalamente | Greg Jefferis Lab | 41,195 |  |
| Remer Tancontian | Mala Murthy Lab, Sebastian Seung Lab | 41,130 |  |
| Nelsie Panes | Mala Murthy Lab, Sebastian Seung Lab | 39,801 |  |
| Laia Serratos Capdevila | Greg Jefferis Lab, Rachel Wilson Lab | 39,798 |  |
| Kendrick Joules Vinson | Mala Murthy Lab, Sebastian Seung Lab | 38,646 |  |
| Anjali Pandey | Greg Jefferis Lab | 37,344 |  |
| Darrel Jay Akiatan | Mala Murthy Lab, Sebastian Seung Lab | 36,987 |  |
| Dustin Garner | Sung Soo Kim Lab | 36,209 |  |
| Ben Silverman | Mala Murthy Lab, Sebastian Seung Lab | 34,676 |  |
| Dharini Sapkal | Greg Jefferis Lab | 31,244 |  |
| Shaina Mae Monungolh | Mala Murthy Lab, Sebastian Seung Lab | 28,721 |  |
| Jay Gager | Mala Murthy Lab, Sebastian Seung Lab | 28,317 |  |
| Krzysztof Kruk | Flyers (citizen scientist) | 27,486 |  |
| Miguel Albero | Mala Murthy Lab, Sebastian Seung Lab | 26,831 |  |
| Katharina Eichler | Greg Jefferis Lab, Seeds Hamepl Lab | 24,481 |  |
| Zeba Vohra | Greg Jefferis Lab | 24,431 |  |
| Emil Kind | Mathias Wernet Lab | 24,052 |  |
| Varun Sane | Greg Jefferis Lab | 23,762 |  |
| annkri (Anne Kristiansen) | Flyers (citizen scientist) | 22,020 |  |
| Chitra Nair | Greg Jefferis Lab | 21,696 |  |
| Márcia dos Santos | Greg Jefferis Lab | 21,209 |  |
| Dhwani Patel | Greg Jefferis Lab | 20,389 |  |
| Imaan F. M. Tamimi | Greg Jefferis Lab | 19,937 |  |
| Michelle Darapan Pantujan | Mala Murthy Lab, Sebastian Seung Lab | 18,846 |  |
| James Hebditch | Mala Murthy Lab, Sebastian Seung Lab | 18,719 |  |
| Alexandre Javier | Greg Jefferis Lab | 17,813 |  |
| Rashmita Rana | Greg Jefferis Lab | 17,547 |  |
| Bhargavi Parmar | Greg Jefferis Lab | 17,133 |  |
| Merlin Moore | Mala Murthy Lab, Sebastian Seung Lab | 16,590 |  |
| Mark Lloyd Pielago | Mala Murthy Lab, Sebastian Seung Lab | 16,566 |  |
| Allien Mae Gogo | Mala Murthy Lab, Sebastian Seung Lab | 16,243 |  |
| Mark Larson | Wei-Chung Lee Lab | 15,692 |  |
| Joseph Hsu | Greg Jefferis Lab, Scott Waddell Lab | 15,496 |  |
| Jacquilyn Laude | Mala Murthy Lab, Sebastian Seung Lab | 14,764 |  |
| Itisha Joshi | Greg Jefferis Lab | 14,717 |  |
| Chereb Martinez | Mala Murthy Lab, Sebastian Seung Lab | 14,638 |  |
| Dhara Kakadiya | Greg Jefferis Lab | 14,544 |  |
| John David Asis | Mala Murthy Lab, Sebastian Seung Lab | 14,202 |  |
| Clyde Angelo Lim | Mala Murthy Lab, Sebastian Seung Lab | 14,139 |  |
| Alvin Josh Mandahay | Mala Murthy Lab, Sebastian Seung Lab | 14,112 |  |
| Thomas Stocks | Flyers (citizen scientist) | 13,683 |  |
| AzureJay (Jaime Skelton) | Flyers (citizen scientist) | 13,540 |  |
| Marchan Manaytay | Mala Murthy Lab, Sebastian Seung Lab | 13,244 |  |
| Kaushik Parmar | Greg Jefferis Lab | 13,136 |  |
| Philipp Schlegel | Greg Jefferis Lab | 12,750 |  |
| Philip Lenard Ampo | Mala Murthy Lab, Sebastian Seung Lab | 12,572 |  |
| Daril Bautista | Mala Murthy Lab, Sebastian Seung Lab | 12,539 |  |
| Irene Salgarella | Greg Jefferis Lab | 12,475 |  |
| John Clyde Saguimpa | Mala Murthy Lab, Sebastian Seung Lab | 11,391 |  |
| Chan Hyuk Kang | Jinseop Kim Lab | 11,129 |  |
| Markus William Pleijzier | Greg Jefferis Lab | 10,498 | Reconstruction of Mushroom Body neurons, Lateral Horn Neurons and Lateral Horn Centrifugal Neurons |
| Marina Gkantia | Greg Jefferis Lab | 10,140 |  |
| Jansen Seguido | Mala Murthy Lab, Sebastian Seung Lab | 9,979 |  |
| Jinmook Kim | Jinseop Kim Lab | 9,879 |  |
| Quinn Vanderbeck | Rachel Wilson Lab | 8,845 |  |
| Cathy Pilapil | Mala Murthy Lab, Sebastian Seung Lab | 8,738 |  |
| Yashvi Patel | Greg Jefferis Lab | 8,485 |  |
| Eva Munnelly | Greg Jefferis Lab | 8,102 |  |
| Olivia Sato | Wei-Chung Lee Lab | 8,055 |  |
| Siqi Fang | Greg Jefferis Lab | 7,981 |  |
| Paul Brooks | Greg Jefferis Lab | 6,838 |  |
| Claire E. McKellar | Mala Murthy Lab, Sebastian Seung Lab | 6,802 |  |
| Christopher Dunne | Greg Jefferis Lab | 6,307 |  |
| Mai Bui | Ken Colodner Lab | 6,228 |  |
| JousterL (Matthew Lichtenberger) | Flyers (citizen scientist) | 5,883 |  |
| edmark tamboboy | Mala Murthy Lab, Sebastian Seung Lab | 5,801 |  |
| Mareike Selcho | Mareike Selcho Lab | 5,565 |  |
| Lucia Kmecova | Seeds Hempel Lab | 5,539 |  |
| Katie Molloy | Wei-Chung Lee Lab | 5,492 |  |
| Alexis E Santana-Cruz | Seeds Hempel Lab | 5,274 |  |
| Janice Salocot | Mala Murthy Lab, Sebastian Seung Lab | 5,133 |  |
| Steven Calle | Seeds Hempel Lab | 4,922 |  |
| Kfay | Flyers (citizen scientist) | 4,886 |  |
| Seongbong Yu | Jinseop Kim Lab | 4,832 |  |
| Arzoo Diwan | Greg Jefferis Lab | 4,787 |  |
| Monika Patel | Greg Jefferis Lab | 4,482 |  |
| Gregory S.X.E. Jefferis | Greg Jefferis Lab | 4,472 |  |
| Sarah Morejohn | Mala Murthy Lab, Sebastian Seung Lab | 4,090 |  |
| Sanna Koskela | Michael Reiser Lab | 3,822 |  |
| bl4ckscor3 (Daniel Lehmann) | Flyers (citizen scientist) | 3,735 |  |
| Celia David | Mala Murthy Lab, Sebastian Seung Lab | 3,611 |  |
| Sangeeta Sisodiya | Greg Jefferis Lab | 3,493 |  |
| Tansy Yang | Janelia | 3,422 |  |
| Selden Koolman | Mala Murthy Lab, Sebastian Seung Lab | 3,384 |  |
| Christa Baker | Mala Murthy Lab | 3,381 |  |
| Szi-chieh Yu | Mala Murthy Lab, Sebastian Seung Lab | 3,376 |  |
| Gerit A. Linneweber | Gerit Linneweber Lab | 3,237 |  |
| Amalia Braun | Alexander Borst Lab | 3,125 |  |
| Sky Cho | Ken Colodner Lab | 2,972 |  |
| Wolf Huetteroth | Wolf Huetteroth Lab | 2,830 |  |
| Brian Reicher | Wei-Chung Lee Lab | 2,794 |  |
| TR77 | Flyers (citizen scientist) | 2,775 |  |
| Marlon Blanquart | Greg Jefferis Lab | 2,662 |  |
| Farzaan Salman | Andrew Dacks Lab | 2,524 |  |
| Hyungjun Choi | Jinseop Kim Lab | 2,373 |  |
| Li Guo | Julie Simpson Lab | 2,095 |  |
| Forrest Collman | Collman | 2,016 |  |
| Marissa Sorek | Mala Murthy Lab, Sebastian Seung Lab | 2,007 |  |
| Joanna Eckhardt | Mala Murthy Lab | 1,995 |  |
| Alisa Poh | Barry Dickson Lab | 1,922 |  |
| Marina Lin | Ken Colodner Lab | 1,920 |  |
| Stefanie Hampel | Seeds Hampel Lab | 1,645 |  |
| Wes Murfin | Citizen scientist | 1,578 |  |
| Peter Gibb | Rachel Wilson Lab | 1,448 |  |
| Zhihao Zheng | Sebastian Seung Lab | 1,421 |  |
| Nidhi Patel | Greg Jefferis Lab | 1,394 |  |
| Lucy Houghton | Sung Soo Kim Lab | 1,357 |  |
| Devon Jones | Mala Murthy Lab, Sebastian Seung Lab | 1,295 |  |
| Alvaro Sanz Diez | Rudy Behnia Lab | 1,284 |  |
| Annalena Oswald | Marion Silies Lab | 1,187 |  |
| Lucas Encarnacion-Rivera | Mala Murthy Lab | 1,164 |  |
| Akanksha Jadia | Greg Jefferis Lab | 1,141 |  |
| Leonie Walter | Mathias Wernet Lab | 1,102 |  |
| Nik Drummond | Alexander Borst Lab | 1,099 |  |
| Xin Zhong | Mathias Wernet Lab | 1,083 |  |
| Benjamin Gorko | Sung Soo Kim Lab | 1,064 |  |
| Fernando J Figueroa Santiago | Seeds Hampel Lab | 1,049 |  |
| István Taisz | Greg Jefferis Lab | 1,043 |  |
| Urja Verma | Greg Jefferis Lab | 1,033 |  |
| Ibrahim Tastekin | Carlos Ribeiro Lab | 1,025 | Tracing taste peg gustatory neurons and downstream neurons |
| Sandeep Kumar | Mala Murthy Lab | 987 |  |
| Yuta Mabuchi | Nilay Yapici Lab | 963 |  |
| Nick Byrne | Wei-Chung Lee Lab | 951 |  |
| Edda Kunze | Gerit Linneweber Lab | 907 |  |
| Thomas Crahan | Sung Soo Kim Lab | 901 |  |
| Hewhoamareismyself (Ryan Margossian) | Flyers (citizen scientist) | 874 |  |
| Iliyan Georgiev | Flyers (citizen scientist) | 825 |  |
| Fabianna Szorenyi | Seeds Hampel Lab | 817 |  |
| Tomke Stuermer | Greg Jefferis Lab | 736 |  |
| Atsuko Adachi | Richard Mann Lab, Rudy Behnia Lab | 695 |  |
| Minsik Yun | Young-Joon Kim Lab | 625 |  |
| Andrearwen | Flyers (citizen scientist) | 607 |  |
| Robert Turnbull | Greg Jefferis Lab | 586 |  |
| Alexander Thomson | Janelia, Michael Reiser Lab | 527 |  |
| a5hm0r | Flyers (citizen scientist) | 516 |  |
| Sebastian Molina-Obando | Marion Silies Lab | 470 |  |
| Connor Laughland | Janelia, Michael Reiser Lab | 469 |  |
| Suchetana B. Dutta | Bassem Hassan Lab | 458 |  |
| Paula Guiomar Alarcón de Antón | Mathias Wernet Lab | 426 |  |
| Patricia Pujols | Seeds Hampel Lab | 423 |  |
| Binglin Huang | Sung Soo Kim Lab | 423 |  |
| Kenneth J. Colodner | Ken Colodner Lab | 421 |  |
| Isabel Haber | Rachel Wilson Lab | 392 |  |
| Albert Lin | Mala Murthy Lab | 362 |  |
| Alexander Shakeel Bates | Greg Jefferis Lab, Rachel Wilson Lab | 340 |  |
| Daniel T. Choe | Jinseop Kim Lab | 340 |  |
| Veronika Lukyanova | Jenny Read Lab | 337 |  |
| Marta Costa | Greg Jefferis Lab | 334 |  |
| Maria Ioannidou | Marion Silies Lab | 332 |  |
| Jonas Chojetzki | Marion Silies Lab | 331 |  |
| Zequan Liu | Xueying "Snow" Wang | 317 |  |
| Haley Croke | Katie von Reyn Lab | 308 |  |
| Gizem Sancer | Mathias Wernet Lab | 308 |  |
| Tatsuo Okubo | Rachel Wilson Lab | 306 |  |
| Miriam A. Flynn | Janelia, Michael Reiser Lab | 297 |  |
| Meghan Laturney | Kristin Scott Lab | 274 |  |
| Benjamin Barger | Salil Bidaye Lab | 273 |  |
| Davi D. Bock | Davi Bock Lab | 255 |  |
| Hyunsoo Yim | Jinseop Kim Lab | 240 |  |
| Anh Duc Le | Denise Garcia Lab | 237 |  |
| Seungyun Yu | Jinseop Kim Lab | 224 |  |
| Yeonju Nam | Jinseop Kim Lab | 221 |  |
| Mavil | Flyers (citizen scientist) | 217 |  |
| Eleni Samara | Alexander Borst Lab | 213 |  |
| Audrey Francis | Gaby Maimon Lab | 196 |  |
| Jesse Gayk | Greg Jefferis Lab | 195 |  |
| Zepeng Yao | Kristin Scott Lab | 194 |  |
| Sommer S. Huntress | Ken Colodner Lab | 192 |  |
| Carolina Manyari-Diaz | Salil Bidaye Lab | 191 |  |
| Raquel Barajas | Carlos Ribeiro Lab | 186 |  |
| Mindy Kim | Wei-Chung Lee Lab | 185 |  |
| Burak Gür | Marion Silies Lab | 182 |  |
| Nils Reinhard | Charlotte Helfrich-Forster Lab | 177 | Tracing of clock and AME neurons |
| Amanda Abusaif | Kristin Scott Lab | 176 |  |
| Anna Li | Rachel Wilson Lab | 173 |  |
| Sven Dorkenwald | Sebastian Seung Lab | 169 |  |
| Fred W Wolf | Fred Wolf Lab | 163 |  |
| Lena Lörsch | Marion Silies Lab | 159 |  |
| Keehyun Park | Jinseop Kim Lab | 155 |  |
| Xinyue Cui | Nilay Yapici Lab | 152 |  |
| Haein Kim | Nilay Yapici Lab | 145 |  |
| Alan Mathew | Greg Jefferis Lab | 141 |  |
| Taewan Kim | Jinseop Kim Lab | 135 |  |
| Guan-ting Wu | National Hualien Senior High School | 124 |  |
| Margarida Brotas | Eugenia Chiappe Lab | 112 |  |
| Cheng-hao Zhang | National Hualien Senior High School | 109 |  |
| Philip K. Shiu | Kristin Scott Lab | 108 |  |
| Shanice Bailey | Greg Jefferis Lab | 102 |  |

**Supplementary Table 2:** Number of annotations by consortium members. Only members with ≥10 annotations are shown.

| Name | Lab Affiliation | total Labels | specific contributions |
| --- | --- | --- | --- |
| Volker Hartenstein | Volker Hartenstein Lab | 13,762 |  |
| Alexander Shakeel Bates | Greg Jefferis Lab, Rachel Wilson Lab | 11,260 |  |
| Krzysztof Kruk | Flyers (citizen scientist) | 11,138 |  |
| Sven Dorkenwald | Sebastian Seung Lab | 6,375 |  |
| Katharina Eichler | Greg Jefferis Lab, Seeds Hampel Lab | 6,366 |  |
| Philipp Schlegel | Greg Jefferis Lab | 3,751 |  |
| David Deutsch | Mala Murthy Lab | 2,549 |  |
| Kaiyu Wang | Barry Dickson Lab | 2,443 |  |
| Yijie Yin | Greg Jefferis Lab | 2,354 |  |
| Stefanie Hampel | Seeds Hampel Lab | 2,129 |  |
| annkri (Anne Kristiansen) | Flyers (citizen scientist) | 1,871 |  |
| Dustin Garner | Sung Soo Kim Lab | 1,782 |  |
| Wolf Huetteroth | Wolf Huetteroth Lab | 1,409 |  |
| AzureJay (Jaime Skelton) | Flyers (citizen scientist) | 1,303 |  |
| Amalia Braun | Alexander Borst Lab | 1,104 |  |
| Austin T Burke | Mala Murthy Lab, Sebastian Seung Lab | 1,013 |  |
| Gizem Sancer | Mathias Wernet Lab | 942 |  |
| Jenna Joroff | Wei-Chung Lee Lab | 900 |  |
| Gregory S.X.E. Jefferis | Greg Jefferis Lab | 716 |  |
| Christa Baker | Mala Murthy Lab | 622 |  |
| Claire E. McKellar | Mala Murthy Lab, Sebastian Seung Lab | 612 |  |
| Markus William Pleijzier | Greg Jefferis Lab | 541 | Reconstruction of Mushroom Body neurons, Lateral Horn Neurons and Lateral Horn Centrifugal Neurons |
| Christopher Dunne | Greg Jefferis Lab | 517 |  |
| Márcia dos Santos | Greg Jefferis Lab | 448 |  |
| Varun Sane | Greg Jefferis Lab | 442 |  |
| Quinn Vanderbeck | Rachel Wilson Lab | 424 |  |
| Lucia Kmecova | Seeds Hampel Lab | 412 |  |
| Steven Calle | Seeds Hampel Lab | 408 |  |
| Rey Adrian Candilada | Mala Murthy Lab, Sebastian Seung Lab | 364 |  |
| Sebastian Molina-Obando | Marion Silies Lab | 347 |  |
| Philip K. Shiu | Kristin Scott Lab | 321 |  |
| Eva Munnelly | Greg Jefferis Lab | 311 |  |
| Remer Tancontian | Mala Murthy Lab, Sebastian Seung Lab | 308 |  |
| Doug Bland | Mala Murthy Lab, Sebastian Seung Lab | 306 |  |
| Ariel Dagohoy | Mala Murthy Lab, Sebastian Seung Lab | 306 |  |
| Joshua Bañez | Mala Murthy Lab, Sebastian Seung Lab | 301 |  |
| Marina Gkantia | Greg Jefferis Lab | 300 |  |
| Jet Ivan Dolorosa | Mala Murthy Lab, Sebastian Seung Lab | 280 |  |
| Nashra Hadjerol | Mala Murthy Lab, Sebastian Seung Lab | 264 |  |
| Zairene Lenizo | Mala Murthy Lab, Sebastian Seung Lab | 236 |  |
| Matt Collie | Rachel Wilson Lab | 223 |  |
| Farzaan Salman | Andrew Dacks Lab | 219 |  |
| Marion Silies | Marion Silies | 183 |  |
| Kendrick Joules Vinson | Mala Murthy Lab, Sebastian Seung Lab | 175 |  |
| John Anthony Ocho | Mala Murthy Lab, Sebastian Seung Lab | 166 |  |
| Thomas Stocks | Flyers (citizen scientist) | 161 |  |
| Kenneth J. Colodner | Ken Colodner Lab | 161 |  |
| Gerit A. Linneweber | Gerit Linneweber Lab | 156 |  |
| Celia David | Mala Murthy Lab, Sebastian Seung Lab | 151 |  |
| TR77 | Flyers (citizen scientist) | 147 |  |
| Megan Wang | Mala Murthy Lab | 130 |  |
| Szi-chieh Yu | Mala Murthy Lab, Sebastian Seung Lab | 129 |  |
| Lucy Houghton | Sung Soo Kim Lab | 123 |  |
| Nils Reinhard | Charlotte Helfrich-Forster Lab | 123 | Identification of clock and AME neurons |
| Ben Silverman | Mala Murthy Lab, Sebastian Seung Lab | 121 |  |
| Regine Salem | Mala Murthy Lab, Sebastian Seung Lab | 111 |  |
| Benjamin Gorko | Sung Soo Kim Lab | 107 |  |
| Nseraf | Flyers (citizen scientist) | 107 |  |
| Mareike Selcho | Mareike Selcho Lab | 102 |  |
| Meet Zandawala | Zandawala Lab | 82 | Identification of peptidergic and gustatory neurons |
| Haein Kim | Nilay Yapici Lab | 72 |  |
| Minsik Yun | Young-Joon Kim Lab | 71 |  |
| Damian Demarest | Michael Pankratz Lab | 70 |  |
| István Taisz | Greg Jefferis Lab | 66 |  |
| Marissa Sorek | Mala Murthy Lab, Sebastian Seung Lab | 62 |  |
| Andrea Sandoval | Kristin Scott Lab | 58 |  |
| Diego A. Pacheco | Mala Murthy Lab | 55 |  |
| Kyle Patrick Willie | Mala Murthy Lab, Sebastian Seung Lab | 53 |  |
| Zhihao Zheng | Sebastian Seung Lab | 51 |  |
| Benjamin Barger | Salil Bidaye Lab | 47 |  |
| Burak Gür | Marion Silies Lab | 44 |  |
| Sandeep Kumar | Mala Murthy Lab | 40 |  |
| Tansy Yang | Janelia | 37 |  |
| Amanda<br>González-Segarra | Kristin Scott Lab | 36 |  |
| Gianna Vitelli | Salil Bidaye Lab | 29 |  |
| Joanna Eckhardt | Mala Murthy Lab | 26 |  |
| Feng Li | Janelia | 26 |  |
| Alvaro Sanz Diez | Rudy Behnia Lab | 24 |  |
| Shuo Cao | David Anderson Lab | 24 |  |
| Haley Croke | Katie von Reyn Lab | 22 |  |
| Nino Mancini | Salil Bidaye Lab | 21 |  |
| Jonas Chojetzki | Marion Silies | 18 |  |
| Gabriella R. Sterne | Gabriella Sterne Lab | 16 |  |
| Kate Maier | Salil Bidaye Lab | 16 |  |
| Amy R Sterling | Sebastian Seung Lab | 15 |  |
| Yuta Mabuchi | Nilay Yapici Lab | 12 |  |
| Lucas<br>Encarnacion-Rivera | Mala Murthy Lab | 10 |  |
| Alexander Del Toro | Kristin Scott Lab | 10 |  |
| Zepeng Yao | Kristin Scott Lab | 10 |  |

